# Hybridization disrupts growth-defense strategies and reveals trade-offs masked in unadmixed populations of a perennial plant

**DOI:** 10.1101/814046

**Authors:** Karl C. Fetter, David M. Nelson, Stephen R. Keller

**Affiliations:** Department of Plant Biology, University of Georgia; Department of Plant Biology, University of Vermont; Appalachian Laboratory, University of Maryland Center for Environmental Science

**Keywords:** Hybridization, trade-offs, growth strategy, *Populus*, *Melampsora*, plant pathology

## Abstract

Organisms are constantly challenged by pathogens and pests which can drive the evolution of growth-defense strategies. Plant stomata are essential for gas-exchange during photosynthesis and conceptually lie at the intersection of the physiological demands of growth and exposure to foliar fungal. Generations of natural selection for locally adapted growth-defense strategies can eliminate variation between traits, potentially masking trade-offs and selection conflicts that may have existed in the past. Hybrid populations offer a unique opportunity to reset the clock on selection and to study potentially maladaptive trait variation before selection removes it. We study the interactions of growth, stomatal, ecopysiological, and disease resistance traits in Poplars *(Populus)* after infection by the leaf rust *Melampsora medusae.* Phenotypes were measured in a common garden and genotyped at 227K SNPs. We isolate the effects of hybridization on trait variance, discover correlations between stomatal, ecophysiology and disease resistance, examine trade-offs and selection conflicts, and explore the evolution of growth-defense strategies potentially mediated by selection for stomatal traits on the upper leaf surface. These results suggest an important role for stomata in determining growth-defense strategies in organisms susceptible to foliar pathogens, and reinforces the contribution of hybridization studies towards our understanding of trait evolution.

## I. INTRODUCTION

Trade-offs arise when selection on fitness is constrained by a negative correlation between a component of fitness and another quantitative trait (Schluter et al., 1991). In plants, an important life history trade-off is between size and the age of first reproduction, with delayed reproduction correlating to longer fertility of the parent and higher quality offspring. However, long-lived organisms like trees must cope with numerous natural enemies, and delaying reproduction can come at a high cost if an organism risks dying before reaching reproductive maturity. Plants have evolved a complex set of constitutive and inducible defenses to cope with their enemies, but they nearly always come at a cost to a component of fitness (Obeso, 2002).

Depending on pathogen prevalence, plant life histories generally evolve towards increased investment in defensive enzymes and compounds paired with slower growth, or decreased investment in defense paired with faster growth (Obeso, 2002). The fast-slow trade-off has been consistently identified in annuals (Tian et al., 2003), perennials (Messina et al., 2002), and long-lived trees (McKown et al., 2014; McKown et al., 2019), although this hypothesis is not without its critics (Kliebenstein, 2016). At the molecular level, the growth-defense trade-off is regulated by cellular signaling and hormonal regulation to direct metabolic activity away from growth and towards defense, or vice versa (Tian et al., 2003; Chandran et al., 2014). Host plant species may evolve a variety of costly defenses to combat disease, including the evolution of structural phenotypes to reduce exposure to pathogens (Gonzales-Vigil et al., 2017), immune systems to detect pathogens (Dangl and Jones, 2001) in order to initiate appropriate responses (Melotto et al., 2006), and resistance via constitutive or inducible synthesis of defensive compounds (Ullah et al., 2018). Hosts may also avoid disease by colonizing new environments where the pathogen is absent (Bruns et al., 2018), or by evolving life history strategies that avoid or compensate for pathogen exposure (Obeso, 2002).

Hybridization is an important mechanism for evolutionary change and has been implicated i multiple phenomena, including the maintenance of species boundaries due to negative selection on advanced generation hybrids (Christe et al., 2016), introgression of beneficial alleles across species barriers (Chhatre et al., 2018), and speciation (Goulet et al., 2017). Phenotypic distributions in hybrid populations often differ from their parental species in important ways. Hybrid vigor, or heterosis, is the enhancement of trait values in early generation hybrids that is a useful tool in crop breeding to increase yields of harvestable organs. Transgressive segregation is a similar concept where trait values in hybrids are elevated or depressed in reference to their parental species (Goulet et al., 2017). Disease resistance will respond to hybridization and can yield undesirable results. For example, hybrid *Salix eriocephala* x *sericea* exhibit a decrease in disease resistance to willow leaf rust *(Melampsora* sp.) by a factor of 3.5 in comparison to their unadmixed parents (Roche and Fritz, 1998). The phenotypic integration of traits that have experienced locally adaptive selection for optimal growth strategies may be disrupted after the arrival of reproductively compatible congeners or genetically divergent demes within the same species that have experienced selection for a different growth strategy. Thus, it is apparent that hybridization can have beneficial and deleterious effects on fitness, but the role of hybridization in disrupting locally adaptive growth strategies and revealing fitness trade-offs that are not evident within the parental species has been little explored.

One scenario where a trade-off between plant growth and defense may arise is the relationship between stomatal traits and infection by fungal pathogens. Stomata are microscopic valves on the surface of the leaf that regulate gas exchange during photosynthesis. Some foliar pathogens enter their hosts via stomata by sensing the topography of the leaf surface for guard cells, and forming an appressorium over a stoma from which a penetration peg grows to invade the mesophyll tissue (Allen et al., 1991). As a dispersal cloud of pathogenic foliar fungal spores moves through an environment containing susceptible hosts, spores will land on the upper and lower leaf surfaces and begin their search for stomata. Physiological models of the benefits and costs to arranging stomata on the upper and/or lower leaf surfaces indicate improved efficiency of transpiration when equal densities of stomata are found on each surface (Muir, 2015). However, leaves may be more prone to infection if the upper leaf surface bears stomata and the risk of pathogen colonization is increased. Here, we see the conditions for the evolution of a growth-defense trade-off mediated by selection on stomata. Hybridization may be expected to shift genotypes along the growth-defense continuum, and when competing growth strategies that rely on various distributions of stomatal architecture traits meet in a hybrid, we may expect a mismatch between optimal growth and defense strategies to yield decreased disease resistance.

In this study, we test for the effects of hybridization on disease resistance, stomatal traits, and growth, and use hybridization as a tool to potentially reveal trade-offs, selection conflicts, and the evolution of different growth strategies between unadmixed and admixed genotypes. We use a sample of naturally formed hybrids between North American poplars infected by the same leaf rust, *Melampsora medusae.* We specifically ask the following questions: 1) how does hybridization change trait distributions, and does it alter their heritable variation; 2) which traits are most predictive of disease resistance; 3) are selection trade-offs and conflicts between growth, disease resistance, and gas exchange observed in hybrids that are not observed in unadmixed populations; and 4) can we identify competing growth-defense strategies in different sets of hybrids potentially fine-tuned by stomatal traits?

## II. MATERIALS & METHODS

### A. STUDY SYSTEM

Poplars *(Populus)* are a genus of predominantly holarctic tree species. Extensive hybridization between species within a section of the genus, as well as between some sections, make the taxonomy of the genus difficult, with some authors identifying 29 to as many as 60 species (DiFazio et al., 2011). Hybrids can be formed from species pairs within and between *Populus* sections *Tacamahaca* and *Aigeiros,* and extensive hybrid zones spontaneously form where the ranges of two reproductively compatible species meet (Suarez-Gonzalez et al., 2018). Western North America contains several well documented hybrid zones (Chhatre et al., 2018; Suarez-Gonzalez et al., 2018), including a tri-hybdrid zone in Alberta, Canada (Floate et al., 2016). The disease we study is from the fungal leaf pathogen *Melampsora medusae* (Fig. 1c,d), a macrocyclic basidiomycete whose aecial host is a larch *(Larix),* and telial host is a poplar. Uredospores (N + N) will emerge from a hyphyal mass of tissue (uredinium) and are able to clonally reproduce on poplar leaves within a single season. The closely related *M. larici-populina* is an agricultural pest that can reduce yields of hybrid poplars grown in agroforestry (Feau et al., 2007).

**Figure 1:**
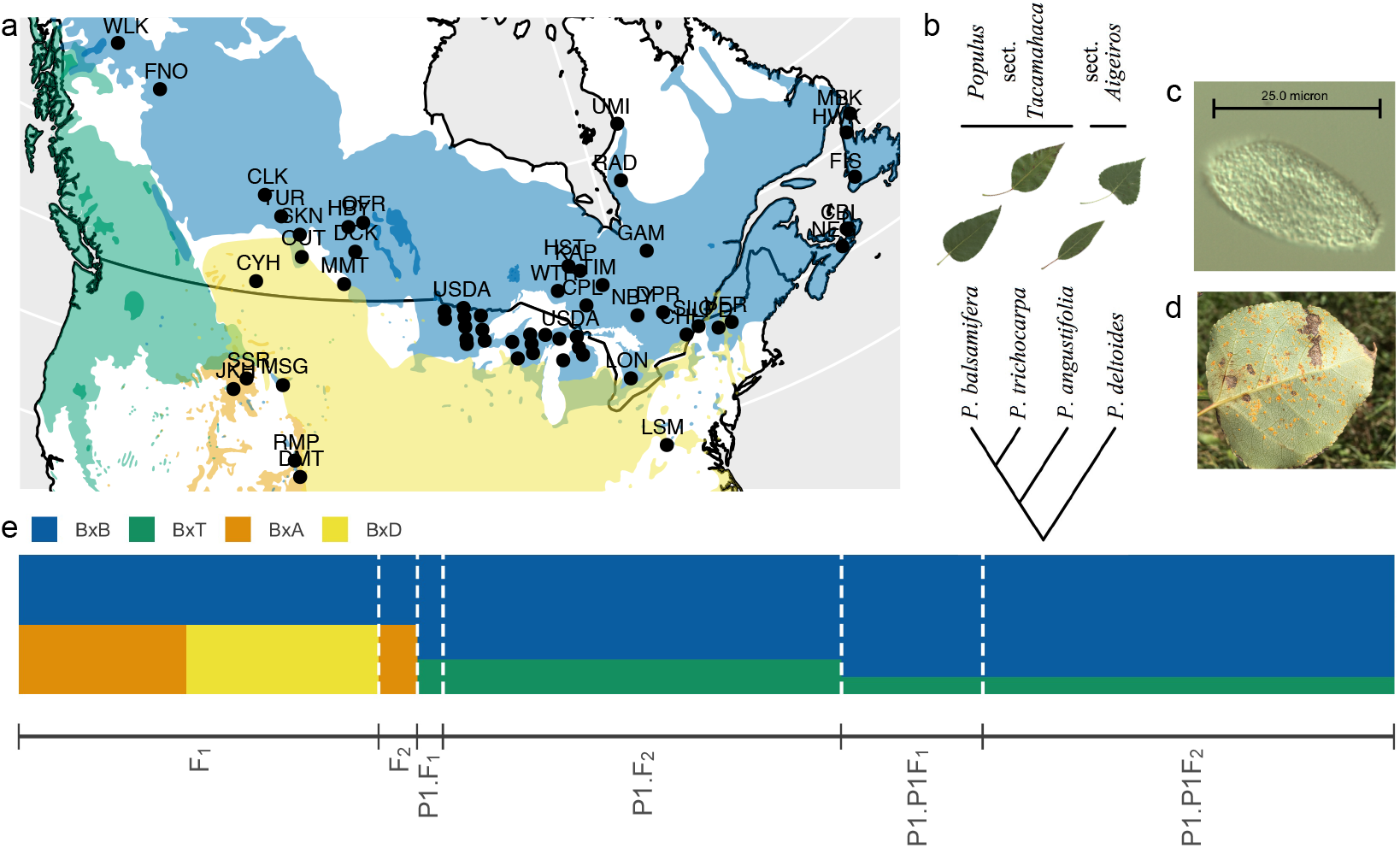
Sample locality map and geographic ranges of four species of poplar: *P. balsamifera* (blue), *P. trichocarpa* (green), *P. angustifolia* (orange), and *P. deltoides* (yellow) (a). See Table S1 for population coordinates and samples sizes. Phylogenetic relationships of the four parental species presented as a cladogram (taxonomy from Eckenwalder, 1996). Representative leaves of each set of poplars from the common garden (b). Uredospore (c) and disease sign (d) of *Melampsora medusae* on the leaf of an *P. balsamifera* x *deltoides* hybrid. The expected proportion of ancestry of hybrid genotypes was calculated from NewHybrid (Anderson and Thompson, 2002) estimates of filial generation (e).

### B. PLANT MATERIALS

In this study, we work with hybrids crossed between a balsam poplar (*P. balsamifera),* and either a black cottonwood *(P. trichocapra),* a narrow-leaf cottonwood (*P. angustifolia),* or an eastern cottonwood (*P. deltoides,* Fig 1b). We lack information on the which species served as the maternal vs. paternal parent for each hybrid, and adopt the convention of listing the *P. balsamifera* parent first. During winter 2013, dormant stem cuttings were collected from 534 trees from 59 populations spanning 9 Canadian provinces and 7 US States (longitudes −55 to −128 °W and latitudes 39 to 60 °N; Fig. 1a, Table S1). The main focus of the 2013 collection was to sample P *balsamifera* cuttings, but as the trees were dormant and some of the diagnostic traits were not immediately visible, a number of putative hybrids were also collected. We collected 32 presumably unadmixed eastern cottonwood (*P. deltoides*) genotypes from central Vermont, USA to serve as a reference population for identifying admixed *P. deltoides* hybrids with population genetic methods. For the 2013 collection, cuttings were grown for one year in a greenhouse, and then planted in the summer of 2014 in a common garden near Burlington, VT (44.444422 °N, −73.190164 °W). Replicates were planted in a randomized design with 2×2 meter spacing and 1,000 ramets were planted. Plants were not fertilized, but were irrigated as-needed during the 2014 growing season to ensure establishment, and then received no supplemental water.

### C. MOLECULAR DATA

Fresh foliage from greenhouse grown plants was used for extracting whole genomic DNA using DNeasy 96 Plant Mini Kits (Qiagen, Valencia, CA, USA). DNA was quantified using a fluorometric assay (Qubit BR, Invitrogen) and confirmed for high molecular weight using 1% agarose gel electrophoresis. We used genotyping-by-sequencing (GBS) (Elshire et al., 2011) to obtain genome-wide polymorphism data for all 534 trees. Genomic sequencing libraries were prepared from 100 ng of genomic DNA per sample digested with EcoT221 followed by ligation of barcoded adapters of varying length from 4-8 bp, following Elshire et al. (2011). Equimolar concentrations of barcoded fragments were pooled and purified with QIAquick PCR purification kit. Purified products were amplified with 18 PCR cycles to append Illumina sequencing primers, cleaned again using a PCR purification kit. The resulting library was screened for fragment size distribution using a Bioanalyzer. Libraries were sequenced at 48 plex (i.e., each library sequenced twice) using an Illumina HiSeq 2500 to generate 100 bp single end reads. Cornell University Institute of Genomic Diversity (Ithaca, NY) performed the library construction and sequencing steps. For the *P. deltoides* reference population, library preparation was performed using the same protocol. DNA sequencing was performed on an Illumina HiSeq 2500 at the Vermont Genetics Network core facility. Raw sequences reads are deposited in NCBI SRA under accession number SRP070954.

We employed the Tassel GBS Pipeline (Glaubitz et al., 2014) to process raw sequence reads and call variants and genotypes. In order to pass the quality control, sequence reads had to have perfect barcode matches, the presence of a restriction site overhang and no undecipherable nucleotides. Filtered reads were trimmed to 64 bp and aligned to the *P. trichocarpa* reference assembly version 3.0 (Tuskan et al., 2006) using the Burrows-Wheeler Aligner (BWA) (Li and Durbin, 2009). Single nucleotide polymorphisms (SNPs) were determined based on aligned positions to the reference, and genotypes called with maximum likelihood in Tassel (Glaubitz et al., 2014). SNP genotype and sequence quality scores were stored in Variant Call Format v4.1 (VCF) files, which were further processed with VCFTools 0.1.11 (Danecek et al., 2011). SNPs with a minor allele frequency < 0.001 were removed, and only biallelic sites were retained. Sites with with a mean depth < 5, genotype quality > 90, and indels were removed. Missing data were imputed with Beagle v5.0 (Browning et al., 2018), and sites with post-imputation genotype probability < 90 and sites with any missingness were removed. After filtering, the final dataset contained 227,607 SNPs for downstream analyses.

Filial generation was estimated separately for BxT and BxD hybrids using NewHybrids (Anderson and Thompson, 2002). Filial generations of *P. balsamifera* x *angustifolia* genotypes were previously estimated by Chhatre et al. (2018). NewHybrids requires reference populations for each species, and *P. trichocarpa* reference genotypes were downloaded from previously published work and 25 genotypes were selected from Pierce Co, Washington from populations known to lack admixture with *P. balsamifera* (Evans et al., 2014). *P. deltoides* reference genotypes were collected from the Winooski and Mad River watersheds in central Vermont. The reference population for *P. balsamifera* was selected from individuals in the SLC, LON, and DPR populations that are known to lack admixture with other *Populus* species (Chhatre et al., 2019). To select loci for distinguishing filial generations, we determined the locus-wise *F_ST_* difference between reference populations. For the BxT analysis, 355 loci with an *F_ST_* difference greater than 0.8 were randomly selected. In the BxD analysis, 385 loci that segregated completely between parental species (i.e. *F_ST_* difference = 1) were randomly sampled. NewHybrids was run using Jeffrey’s prior for π and *θ* for 200,000 sweeps with 100,000 discarded as burn-in. The expected proportion of the genome from each parental species was calculated as:

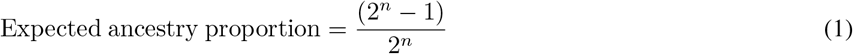

where n = the number crossing events. The admixture status (i.e. admixed/unadmixed) and hybrid set (i.e. BxB, BxT, BxA, or BxD) was determined from the expected ancestry proportions.

### D. TRAIT DATA

All traits were measured from common garden grown trees in 2015, and disease severity was measured again in 2016 (Table 1).

**Table 1:** Trait definitions, abbreviations, and units.

| Definition | Abbvr. | Units |
| --- | --- | --- |
| Disease |  |  |
| Disease Resistance 2015 | $R_1$ | ordinal |
| Disease Resistance 2016 | $R_2$ | ordinal |
| Stomatal patterning |  |  |
| Upper stomatal density | $D_U$ | $\text{mm}^{-2}$ |
| Lower stomatal density | $D_L$ | $\text{mm}^{-2}$ |
| Upper stomatal size | $S_U$ | $\mu\text{m}^2$ |
| Lower stomatal size | $S_L$ | $\mu\text{m}^2$ |
| Lower stomatal cover | $f_{S_U}$ | none |
| Upper stomatal cover | $f_{S_L}$ | none |
| Total stomatal cover | $f_S$ | none |
| Upper interstomatal distance | $U_U$ | $\mu\text{m}$ |
| Lower interstomatal distance | $U_L$ | $\mu\text{m}$ |
| Upper anatomical maximum stomatal conductance | $g_{s,\max_U}$ | $\text{mol m}^{-2}\text{s}^{-1}$ |
| Lower anatomical maximum stomatal conductance | $g_{s,\max_L}$ | $\text{mol m}^{-2}\text{s}^{-1}$ |
| Total anatomical maximum stomatal conductance | $g_{s,\max}$ | $\text{mol m}^{-2}\text{s}^{-1}$ |
| Stomatal density ratio | SR | none |
| Stomatal area ratio | AR | none |
| Stomatal cover ratio | $f_S R$ | none |
| Ecophysiology |  |  |
| Relative growth rate | $G$ | cm |
| Carbon:Nitrogen | CN | none |
| Leaf percent carbon | %C | % |
| Leaf percent nitrogen | %N | % |
| Carbon isotope discrimination | $\Delta^{13}\text{C}$ | ‰ |
| Nitrogen isotope value | $\delta^{15}\text{N}$ | ‰ |
| Specific leaf area | SLA | $\text{mm}^2 \text{mg}^{-1}$ |
| Chlorophyll content index | CCI | none |

To control for trait variation between leaves due to age or environmental effects (e.g. aspect, or light), the first fully expanded leaf on the dominant shoot was sampled. Stomatal patterning and ecophysiology traits were measured from the same leaf. The severity of naturally inoculated leaf rust disease was phenotyped in both years using an ordinal scale from zero to four created by LaMantia et al. (2013) where: 0 = no uredinia visible, 1 = less than five uredinia per leaf on less than five leaves, 2 = less than five uredinia per leaf on more than five leaves, 3 = more than five uredinia per leaf on more than five leaves, and 4 = more than five uredinia on all leaves. Disease severity was converted to resistance (*R*) with the function *R* = −1 x Severity + 6. A mature larch tree (approx. 20m tall) was located approximately 100 m from the garden site, and we assume a uniform distribution of aeciospore inoculum into the garden. Using microscopy, the pathogen was visually confirmed as *M. meduscae* by the ellipsoid to obovoid shape of uredospores, the size range (mean = 28.3 μm, min = 19.6 μm, max = 34.45 μm, N = 17), and the presence of a smooth equatorial region on the spore flanked by polar regions with papillae (Van Kraayenoord et al., 1974) (Fig. 1c).

To collect isotopic, elemental, and specific leaf area (SLA) data, three hole punches (diameter = 3 mm) were sampled in June 2015 from a central portion of each leaf adjacent to, but avoiding the central leaf vein. Hole punches were dried at 65 °C to constant mass. Approximately 2 mg of foliar tissue from each sample was weighed into a tin capsule and analyzed for %C, %N, δ^13^C, and δ^15^N using a Carlo Erba NC2500 elemental analyzer (CE Instruments, Milano, Italy) interfaced with a ThermoFinnigan Delta V+ isotope ratio mass spectrometer (Bremen, Germany) at the Central Appalachians Stable Isotope Facility (CASIF) at the Appalachian Laboratory (Frostburg, Maryland, USA). %C and %N were calculated using a size series of atropine. The δ^13^C and δ^15^N data were normalized to the VPDB and AIR scales, respectively, using a two-point normalization curve with laboratory standards calibrated against USGS40 and USGS41. The long-term precision of an internal leaf standard analyzed alongside samples was 0.28%o for δ^13^C and 0.24%o for δ^15^ N. Isotopic results are reported in units of per mil (‰). Carbon isotope discrimination against ^13^C (Δ^13^C) was calculated according to Farquhar et al. (1982) as:

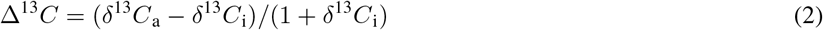

where the δ^13^C_a_ value (−8.456 ‰) was the mean value in 2015 measured at NOAA Mauna Loa observatory (White et al., 2011). Higher Δ^13^C values indicate increased intracellular (C_i_) relative to atmospheric (C_a_) CO_2_ concentrations as a result of greater stomatal conductance and/or lower photosynthetic assimilation rates in C_3_ plants. Δ^13^C is a useful metric of intrinsic water-use efficiency (WUE), or the ratio of photosynthesis to stomatal conductance, since both are influenced by C_i_/C_a_. To calculate SLA, or the ratio of fresh punch area to dry leaf mass, three oven dried hole punches per leaf were massed collectively. Relative growth rate (G) was measured as the height increment gain (cm) between the apical bud in 2015 and the previous year’s bud scar on the most dominant stem. The chlorophyll content index (CCI) was measured with a Konica Minolta SPAD 502 (Konica Minolta Sensing Americas, Inc) and the average of three measurements from the central portion of a leaf was recorded.

Stomata patterning traits were measured from micrographs of nail polish casts of the lower (abaxial) and upper (adaxial) leaf surfaces. Leaves were collected from the field and placed in a cooler until processed in the lab. Nail polish peels were made and mounted on slides without a cover slip. Two non-overlapping areas without large veins from each peel were imaged (N = 1894) with an Olympus BX-60 microscope using differential interference contrast. Stomata density (D) was estimated using the machine learning protocol of StomataCounter (Fetter et al., 2019). To verify the automatic counts, stomata were manually annotated on each image using the image annotation tool. The correlation between automatic and manual count was *r* = 0.99, automatic counts were used. Stomatal aperture pore length was measured from micrographs in ImageJ (Schneider et al., 2012) by overlaying four equally spaced lines across an image, and then measuring a single aperture pore from each segmented region for a total of five observations per image. Stomatal size (*S*), stomatal cover (*f_S_*), defined as the covering fraction of the leaf surface by stomatal aperture pores, and theoretical maximum gas exchange (*g*_s,max_), were respectively calculated as

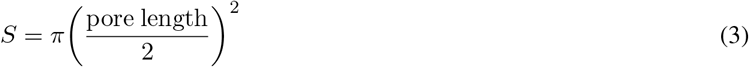

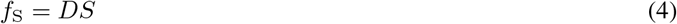

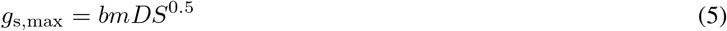

where *b* is the diffusion of coefficient of water vapor in air (*b* = 0.001111607) and *m* is a morphological constraint of stomatal guard cell length and width, aperture pore length and depth (*m* = 0.4320532; see Sack and Buckley, 2016 for details). Interstomatal distances (*U*) were calculated separately for upper and lower leaf surfaces as

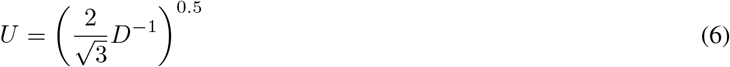

following Muir (2020). Ratios of stomatal density (SR), stomatal area (AR, calculated from size), and the stomatal cover ratio (*f*_S_R), are calculated as a ratio of the upper leaf surface trait to the total. For example,

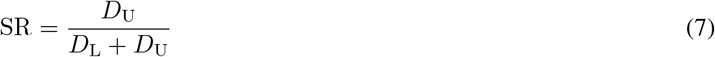

Using this formula, a SR is bound by 0 and 1, and a value of 0.5 indicates an equal density of stomata on the upper and lower leaf surfaces.

### E. QUANTITATIVE GENETIC ANALYSES

To investigate how hybridization changes trait distributions and heritable variation, we fitted a series of mixed-effects models with factors describing different levels of hybrdization, fitted a partial-least squares (PLS) model, and estimated heritability. The hybridization-level models were fit with brms (Bürkner, 2017) using unscaled trait data, and then again to rescaled trait data with a mean of zero and variance of two-times the standard deviation *(sensu* Gelman, 2008). In total, six sets of models were fit, substituting a different vector of hybiridization level in in each model, given by

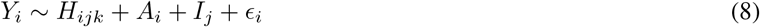

where *Y* is a trait, *H* was a vector of 1) admiuxture status (i.e. admixed or unadmixed); 2) hybrid set (i.e. BxB, BxT, BxA, or BxD); or 3) the filial generation (i.e. *F*_1_, *F*_2_, etc.); *A* is a matrix of xy garden position coordinates, *I* is the random effect of the individual’s genotype, e is the error term, and *i,j, k* represent the levels of replicate, individual, and the hybridization vector, respectively. Each model was run using four Markov chains, with 4000 burn-in and 8000 sampling iterations. Model mixing was improved by setting the max_treedepth to 15 and adapt_delta to 0.99. For each trait, various relevant distribution models were fit to the data and the best family chosen using using leave-one-out (LOO) cross validation and posterior predictive checks (Table S3). A PLS model was fit with the package mixOmics (Rohart et al., 2017) using canonical correlation, where the X matrix was a column vector of the expected ancestry proportions, and the Y matrix was a column vector of traits. Broad-sense heritability *(H*^2^) was estimated separately for unadmxied and admixed data sets using brms. Models where fit with garden position and genet identity using the same model-run specifications and distribution family selection methodology described above. H^2^ was estimated as,

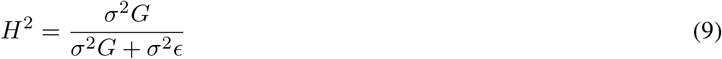

where *σ*^2^*G* represented the variance attributable to genotype and *σ*^2^*ϵ* represented the environmental and error variances. The posterior median and 90% credible intervals were recorded. The *RV* coefficient between the absolute value of the centered PLS scalar product matrix and the centered heritability matrix was estimated with FactoMineR (Lê et al., 2008) and a p-value estimated with permutation testing.

Covariation of predictors to disease resistance was investigated by fitting four multi-level, multi-response models in brms with a cumulative family distribution given by

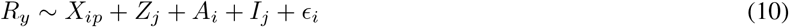

where *R_y_* is *R*_1_ or *R*_2_, *X_ip_* is a matrix of *p* predictors measured from each replicate, *Z* is a vector of the expected ancestry proportion of P *balsamifera* from each genotype, and *A* and *I* are defined as above. Some stomatal patterning traits were linear combinations of *D* and *S* and were removed, including *U*_U_, *U*_L_, *f*_S_L__, *f*_S_U__, *g*_s,max_U__, and *g*_s,max_U__. Variance from %C and %N were included in the model as their ratio, CN. The four models differed in their inclusion of stomatal patterning and stomatal ratio traits (Table S4). Model-run parameters were the same as above, and fitted models were evaluated with LOO cross validation.

To investigate the covariance between growth, disease resistance, *g*_s,max_, and how admixture can reveal trade-offs and selection conflicts, we first estimated marginal BLUPs (mBLUPs) for the three traits, rescaled the data (*μ* = 0, *σ*^2^ = 2 * *σ*), and fit an interaction model in brms separately for *R*_1_ and *R*_2_. mBLUPs were fit by modeling the trait as a function of garden xy coordinates and the random effects of individual, and then adding the intercept to each random effect. The interaction model was given by

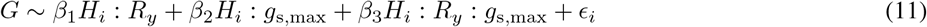

where *H* is a vector of factors describing the hybrid set (i.e. BxB, BxT, BxA, or BxD), *R_y_* is defined as above, and *β_N_* are the regression coefficients. We used a gaussian distribution family and the model-run parameters previously described. Evidence for trade-offs between traits was considered present if the product of the slopes was negative; similarly, selection conflicts were inferred if the product of the slopes from a trade-off were negative (Schluter et al., 1991). Parameter estimates with 95% credible intervals (CI) that do not overlap zero were considered significant. A path analysis was performed to infer the effect of *g*_s,max_ on growth by summing its independent effects estimated from the regression coefficients, calculated as *R*_g_s,max,G__ = *β*_2_ + (*β*_3_ * *β*_1_).

Finally, we further explored the resistance-trait co-variance (Eq. 10) and trade-off models (Eq. 11) by fitting a model to rescaled data to search for contrasting growth-defense trait syndromes that are mediated by selection for different values of gs,max or *D*_U_, given by

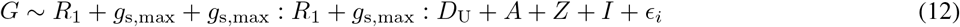

where abbreviations are given in Table 1 and model parameters the same as above. We chose to fit the model to *R*_1_, as there were stronger support for trade-offs than in *R*_2_.

## III. RESULTS

### A. TRAIT DISTRIBUTIONS AND HYBRIDIZATION EFFECTS

Extensive hybridization and backcrossing was revealed from the NewHybrids analyses (Fig 1e, Table 2, Tables S2a, S2b, S2c). Our sample collection protocol was designed to target non-hybrid genotypes, thus the distribution of hybrids in our sample is less than what is expected on the landscape. Nevertheless, we observed hybrids with P *deltoides* at the F_1_ generation, with *P. angustifolia* at the F_1_ and F_2_ generations, and with P *trichocarpa* at advanced stages of backcrossing into P *balsamifera.*

**Table 2:**
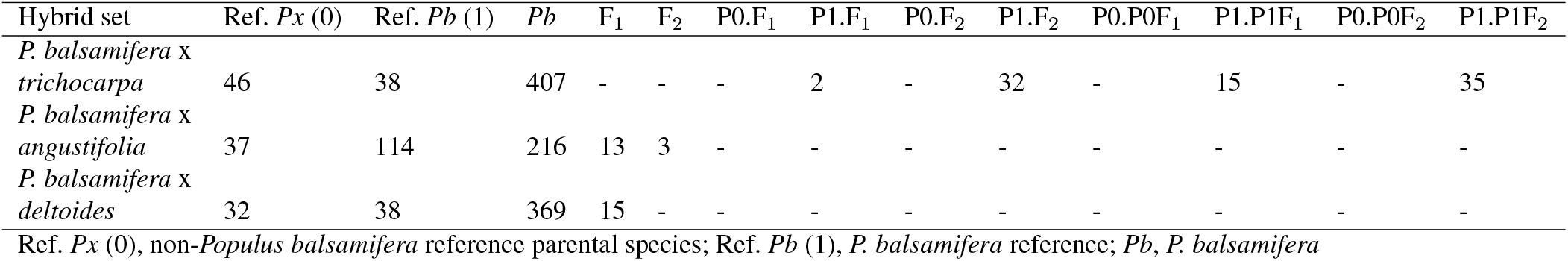
Sample sizes of reference populations from which segregating loci were selected and NewHybrids estimates for the number of genotypes within each filial generation.

| Hybrid set | Ref. $P_x$ (0) | Ref. $P_b$ (1) | $P_b$ | $F_1$ | $F_2$ | $P0.F_1$ | $P1.F_1$ | $P0.F_2$ | $P1.F_2$ | $P0.P0F_1$ | $P1.P1F_1$ | $P0.P0F_2$ | $P1.P1F_2$ |
| --- | --- | --- | --- | --- | --- | --- | --- | --- | --- | --- | --- | --- | --- |
| *P. balsamifera* x *trichocarpa* | 46 | 38 | 407 | - | - | - | 2 | - | 32 | - | 15 | - | 35 |
| *P. balsamifera* x *angustifolia* | 37 | 114 | 216 | 13 | 3 | - | - | - | - | - | - | - | - |
| *P. balsamifera* x *deltoides* | 32 | 38 | 369 | 15 | - | - | - | - | - | - | - | - | - |

Ref. $P_x$ (0), non-*Populus balsamifera* reference parental species; Ref. $P_b$ (1), *P. balsamifera* reference; $P_b$ , *P. balsamifera*

Disease resistance was measured during two years and showed a left skewed distribution with the majority of genotypes resistant to *M. medusae* in both *R*_1_ (55%) and *R*_2_ (80%) (Fig. S1). Zero-inflated distributions were observed for upper leaf surface stomatal patterning and ratio traits, while stomatal patterning of the lower leaf surface traits were gaussian distributed. AR was zero-inflated and the non-zero values in the tail had a median of 0.43 with a minimum value of 0.29 and a maximum of 0.54. Similarly, SR was zero-inflated and the tail had a median of 0.18 with a minimum of 0.05 and a maximum of 0.44. Admixed genotypes, along with 43 of 315 unadmixed BxB individuals, had positive stomatal ratio values. Ecophysiolgy traits were generally gaussian distributed. We observed high co-variance of stomatal patterning traits to eachother, as well as moderate to high correlations between resistance and ecophysiology traits (Fig. S2).

Hybridization had a considerable effect on the distribution of trait values (Eq 8) measured at the level admixture status (Fig. S3), hybrid set (Fig. S4), and filial generation (Fig. S5). Distribution families for unscaled and rescaled data were typically the same for a given trait, but varied considerably across traits (Table S3). Admixture status influences resistance in both years. Unadmixed genotypes have the highest probability of exhibiting complete resistance in both *R*1 (*P* = 0.79) and *R*2 (*P* = 0.97), while for admixed genotypes, the lowest resistance score had the highest probability in *R*_1_ (P = 0.34, Fig. S3). In 2016, admixed genotypes exhibited increasing probability of resistance from 1 (least resistance) to 5 (completely resistance) (Fig. S3). By rescaling and centering traits before fitting models, we can plot the conditional effects of hybridization jointly and observe shifts in integrated sets of traits. Viewing hybridization at its most fundamental level, whether a genotype is admixed or not, we observed a coordinated increase of upper stomatal patterning traits, a decrease in disease resistance and growth, while ecophysiology traits remained largely unchanged (Fig. S6a). When considering different sets of hybrids the deviation of the conditional effects from zero increased from unadmixed BxB genotypes to BxD hybrids (Fig. 2a). Upper stomatal patterning traits diverged first in BxT hybrids and remained elevated in BxA hybrids, but some decreased in BxD hybrids, including *D*_U_. In BxB, BxT, and BxA hybrids *S*_L_, *f*_S_L__ and *f*_S_ remained at similar values and increased in BxD hybrids. *D*_L_ is highest in BxB and decreased in each subsequent hybrid set. Resistance decreased with increasing phylogenetic distance of the non- *balsamifera* parent. Growth decreased in BxT and BxA hybrids relative to BxB, and was elevated in BxD genotypes, of which all are F_1_ generation hybrids (Fig. 2b). Δ^13^C, frequently interpreted as a measure of WUE, was elevated for BxA hybrids, indicating decreased WUE in these hybrids. The remaining ecophysiology traits remain largely unchanged between hybrid sets (Figs. 2a, S4). Although we had limited ability to estimate variance components for parameters in the filial generations of F2 (N=3) and P1.F_1_ (N=2), we can generally report that trait variation was largest at the F_2_ and F_1_ generations, and remains high until the P1.P1F_1_ generation, and is lowest in the unadmixed P *balsamifera* (Fig. S6b). Resistance to *M. medusae* was highest in unadmixed genotypes, P1.P1F_2_, and P1.P1F_1_ filial generations. After P1.F_2_, genotypes had decreased resistance. Stomatal patterning traits were generally lower in the unadmixed filial generation, and increased in value and in variance in subsequent filial generations. After the P1.F_2_ generation, SR, AR, *f*_S_R, *D*_u_, *f*_S_, *f*_S_U__, became elevated and remained so.

**Figure 2:**
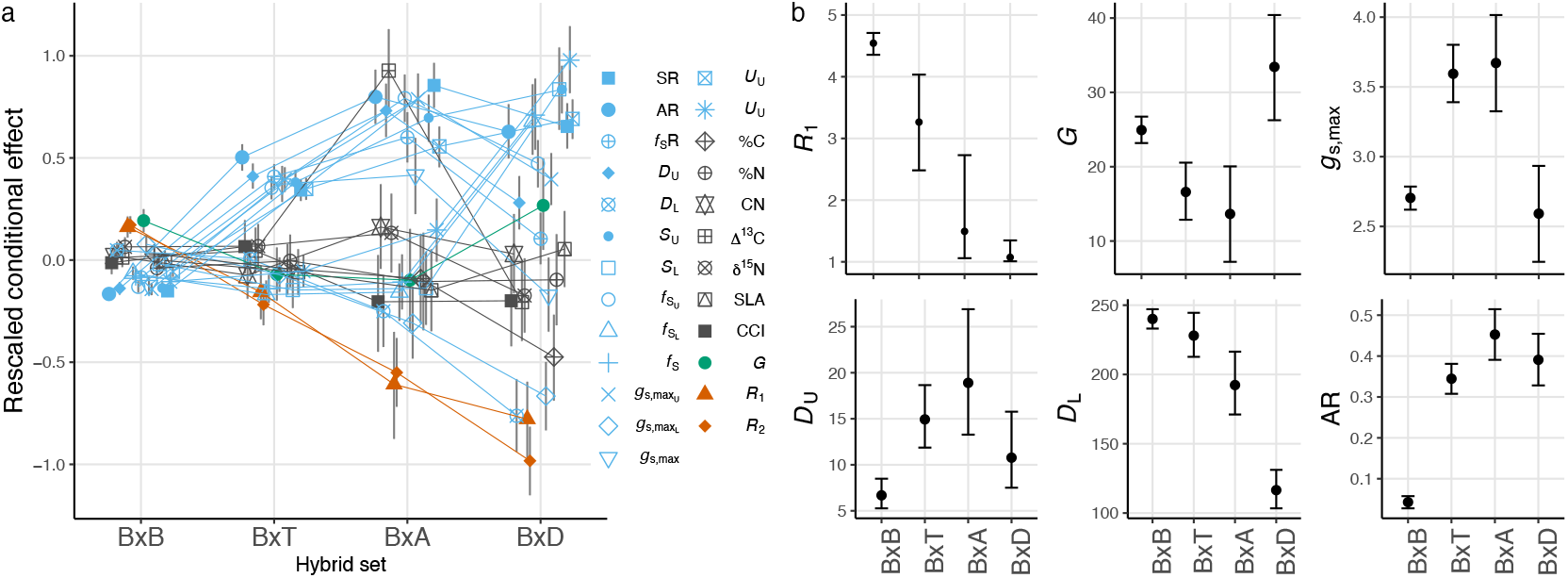
Point estimates and 95% credible intervals for the conditional effects of hybridization on trait values. Rescaled and centered trait data (a) and unscaled data (b) were analyzed with mixed-effects models to observe the effects of hybrid set (Eq. 8). See Table 1 for trait abbreviations and Table S3 for distribution families of each model.

*H*^2^ estimates ranged widely from 0.01 to 0.8 (Fig. 3a). *H*^2^ increased from 0.38 to 0.45 as a result of admixture (t-test: t = 1.0815, df = 47.62, p-value = 0.2849, N = 50, 25 per set). For both the admixed (red) and unadmixed (black) data sets, the upper stomatal traits had a mean *H*^2^ estimate of 0.59, the lower stomatal traits 0.52, and the stomatal ratio traits had a mean of 0.56. In both years, *H*^2^ estimates for disease resistance were higher in the admixed data set and lower in the unadmixed. Including *G*, the ecophysiology traits had a mean H^2^ of 0.2. PLS analyses were conducted to investigate the effect of changes in the expected ancestry proportion from each species on the traits. The scalar product between pairs of vectors in the X and Y matrices indicate the degree of correlation between variables. Using hierarchical clustering of the scalar products, we observed four blocks of traits containing: resistance, *D*_L_, g_s,max_L__, and %C (block 1); ecophysiology traits, growth, and *U*_U_ (block 2); *S*_L_ *U*_L_, *f*_S_L__ and *f*_S_ (block 3); and *g*_s,max_, SR, AR, *f*_s_R, and the upper stomatal patterning traits (block 4). Block 3 and 4 contained, in general, the stomatal traits, with some lower stomatal traits clustering into block 3 and the ratio and upper stomatal traits in block 4. Overall the scalar products were positively correlated to increasing *P. balsamifera* ancestry in block 1 and negatively correlated in block 4. Block 2 has scalar products that were neither positive nor strongly negative, while block 3 is largely characterized by positive correlation to *P. deltoides* ancestry (Fig. 3b). The correlation of the absolute value of the mean-centered scalar products to *H*^2^ was moderate *(RV* = 0.35, p-value < 0.01).

**Figure 3:**
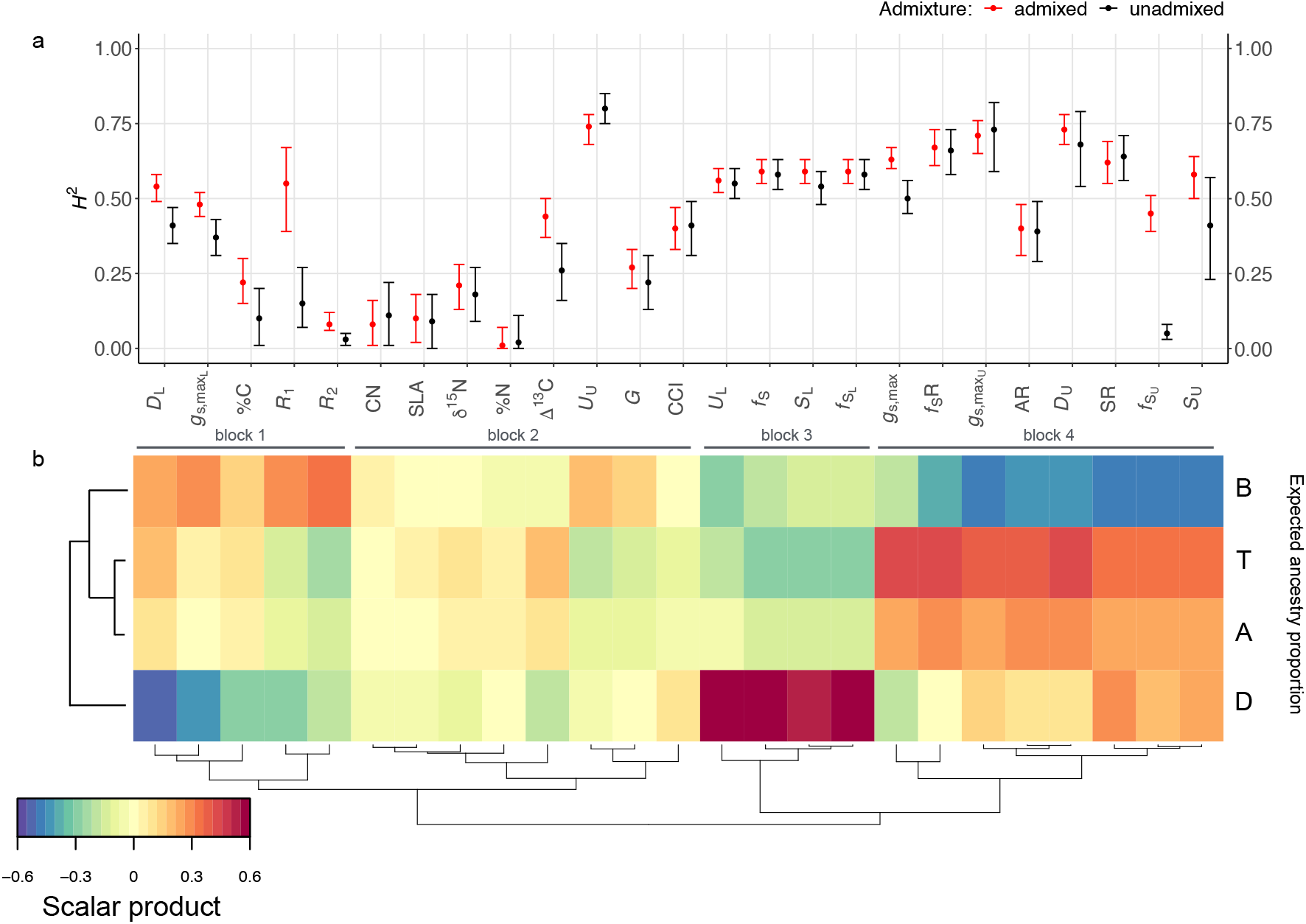
Median broad-sense heritability (*H*^2^) and 90% credible intervals estimated from data sets containing both admixed and unadmixed samples (red) or only unadmixed samples (black) (a). Partial least squares (PLS) regression of ancestry proportions (X matrix) and traits (Y matrix) (b). B, T, A, and D represent vectors of the expected ancestry proportions for each individual estimated by NewHybrids (Table 2). Correlations between pairs of traits are indicated by the color ramp, which represents the product of variable loading vectors U’ and V’ from the X and Y matrices, respectively. The *RV* coefficient between the scalar product and *H*^2^ matrices is 0.35 (p-value < 0.01). See Table S2 for *H*^2^ estimates.

### B. MULTI-RESPONSE REGRESSION OF DISEASE RESISTANCE

A multi-response model was fit simultaneously for *R*_1_ and *R*_2_ to a subset of the stomatal patterning, growth, ecophysiology, and ancestry data (Eq. 10, Fig. 4). The model was fit on data collected at the replicate level, allowing us to include experimental design effects, and to account for individual-level variation with a random effect of genotype. We used the difference of the expected log point-wise predictive density (ΔELPD) to rank models, and the model which included upper and lower *D* and *S* variables seperately, but excluded *f*_S_R was favored (Table S4). The proportion of *P. balsamifera* ancestry explained the most variance in the model and was positively correlated to the disease resistance responses (regression coefficients: *R*_1_ = 15.6; *R*_2_ = 39.9). In *R*_1_ at the 95% CI, *D*_U_, AR, *G*, Δ^13^C were negatively correlated, while CN was positively correlated. At the 66% CI, *f*_S_ and CCI were negatively correlated, while *g*_s,max_ and SR were positively correlated. In *R*_2_ at the 95% CI, *D*_L_, *S*_U_, *δ*^15^N were negatively correlated, while CN was positively correlated. At the 66% CI, *S*_L_ SLA, and CCI were negatively correlated, while *f*_S_, *g*_s,max_ were positively correlated (Fig. 4A).

**Figure 4:**
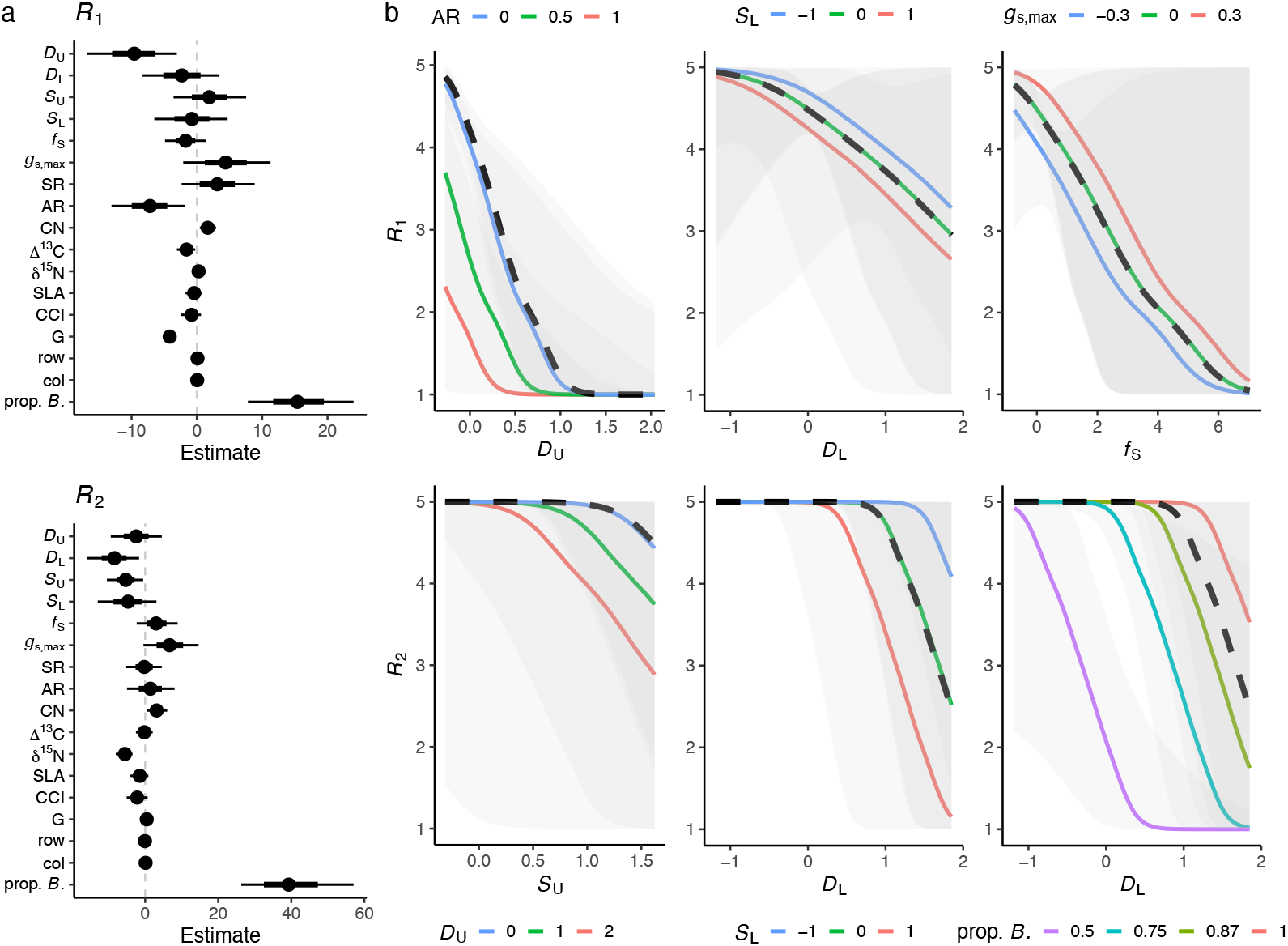
Multiple regression of stomatal patterning and ecophysiology traits on disease resistance within a Bayesian multi-response model (Eq. 10). Median regression coefficient estimates and their 66% and 95% quantile credible intervals (a). Predicted conditional effect of the independent variable (dashed line) and interactions terms (solid lines) are given in the right panels (b).

Through plotting interactions of traits against *R*_1_ and *R*_2_, we explored the effect of a third trait while controlling for an independent variable (Fig. 4B). After accounting for the negative correlation between *D*_U_, increasing AR (i.e. shifting stomatal area to the upper surface) decreases resistance for genotypes with low *D*_U_. At a given level of *D*_L_, increasing *S*_L_ decreases resistance. *f*_S_ is negatively correlated with *R*_1_, and increasing *g*_s,max_ at a given level of *f*_S_ increases resistance. In *R*_2_, *S*_U_ is negatively correlated with resistance at high values, but resistance is lost even faster when the density of stomata on the upper surface increases. A similar pattern is observed on the lower leaf surface. Finally, increases in the proportion of *P. balsamifera* ancestry ameliorated the negative effects of increases in *D*_L_.

### C. SELECTION TRADE-OFFS, CONFLICTS, AND GROWTH-STRATEGIES

To determine if there was evidence for evolutionary trade-offs in unadmixed and hybrid populations, and if trade-offs were revealed in some hybrid sets but not others, multiple regression models with two-way and three-way interaction terms were fit (Eq. 11). A significant trade-off between *R*_1_ and G was observed in BxB, BxT, and BxD hybrid sets, but not in BxA hybrids, which had a positive correlation (Table 3). Increasing values of *g*_s,max_ had a significant trade-off to G in BxT hybrids, but a significant positive effect in BxD hybrids. In BxB genotypes, increasing *g*_s,max_ had a significant positive effect on G through it’s independent effects on *R*_1_, as observed in the three-way interaction (*β*_3_). Fewer interaction terms were significant in the regressions using *R*_2_, but a significant trade-off between G and *R*_2_ was observed in BxD hybrids, and the slope was reversed in BxA hybrids.

**Table 3:** Trade-offs, selection conflicts, and path analysis (*R*_*g*_s,max_,*G*_) of the effects of theoretical maximum stomatal conductance (*g*_s,max_) and disease resistance (*R_y_*) on growth (*G*). Coefficients were estimated with hybrid set (*H*) as an interaction term (Eg. 11). Negative regression coefficients or their products indicate trade-offs or selection conflicts, respectively. The path analysis sums each independent path of the effect of *g*_s,max_ on *G*, as indicated by the path diagram. Parameter estimates that do not overlap zero at a 95 % CI indicated by bold.

| $R_1$ | | | | | |
| --- | --- | --- | --- | --- | --- |
| Hybrid set | $H : R_1 (\beta_1)$ | $H : g_{s,\max} (\beta_2)$ | $H : g_{s,\max} : R_1 (\beta_3)$ | $\beta_1 \times \beta_2$ | $R_{g_{s,\max},G}$ |
| BxB | **-0.76** | 0.03 | **0.17** | -0.03 | -0.22 |
| BxT | **-0.28** | **-0.35** | 0.18 | 0.08 | **-0.4** |
| BxA | **0.71** | -0.12 | -0.57 | -0.07 | -0.53 |
| BxD | **-0.21** | **0.55** | -0.03 | -0.04 | 0.55 |

| $R_2$ | | | | | |
| --- | --- | --- | --- | --- | --- |
| Hybrid set | $H : R_2 (\beta_1)$ | $H : g_{s,\max} (\beta_2)$ | $H : g_{s,\max} : R_2 (\beta_3)$ | $\beta_1 \times \beta_2$ | $R_{g_{s,\max},G}$ |
| BxB | -0.06 | 0.09 | -0.02 | 0 | 0.09 |
| BxT | 0.2 | -0.29 | -0.18 | -0.04 | -0.36 |
| BxA | **0.5** | -0.15 | -0.4 | -0.05 | -0.41 |
| BxD | **-0.3** | -1.3 | -1.6 | 0.27 | -0.75 |

None of the selection conflicts between resistance and *G* or *g*_s,max_ and *G* were statistically significant; nevertheless, selection conflicts were observed for BxB, BxA, and BxD hybrid sets in *R*_1_, and for BxT and BxA sets in *R*_2_. The path analysis investigated the cumulative effect of *g*_s,max_ on *G* through two independent pathways. Path analysis results varied by year and by hybrid set. In *R*_1_, the path analysis yielded a significant negative result for BxT, and non-significant negative results for BxB and BxA hybrid sets. In *R*_2_, none of the paths were signficant, but the sign of the path reversed in BxB and BxD hybrid sets (Table 3).

Finally, we explored the data for the presence of contrasting growth-resistance strategies, possibly fine-tuned by variation of *g*_s,max_ and *D*_U_ (Eq 12). We only explored variation of *R*_1_, as the relationships between traits were stronger in the 2015 data (Table 3). We again recovered the negative relationship between resistance and growth (regression coefficient = −0.59, significant at 95% CI, Fig 5a). BxB genotypes anchored the low-growth/high-resistance growth strategy and BxT genotypes were intermediate to BxA and BxD genotypes which had less resistance. The growth of non-*P. balsamifera* accessions is possibly impacted by the disease, which could account for growth that is substantially less than the predicted values from the model. Overall, *g*_s,max_ had a positive slope with G (regression coefficient = 0.13, significant at 95% CI), and two contrasting strategies for *g*_s,max_ were observed which were fine tuned by *D*_U_ (*g*_s,max_: *D*_U_ regression coefficient = 0.24, significant at 95% CI). The negative effects of high resistance on *G* can be ameliorated by decreasing *g*_s,max_ (*R*_1_: *g*_s,max_ coefficient = −0.1 significant at 66% CI). At high values of *g*_s,max_, growth can be increased with higher values of *D*_U_; while at low values of *g*_s,max_, higher growth is achieved with lower values of *D*_U_. The low *g*_s,max_-*D*_U_ growth strategy is occupied by BxB genotypes and BxD hybrids, while BxA and BxT hybrids occupy the alternative strategy (Fig. 5b).

**Figure 5:**
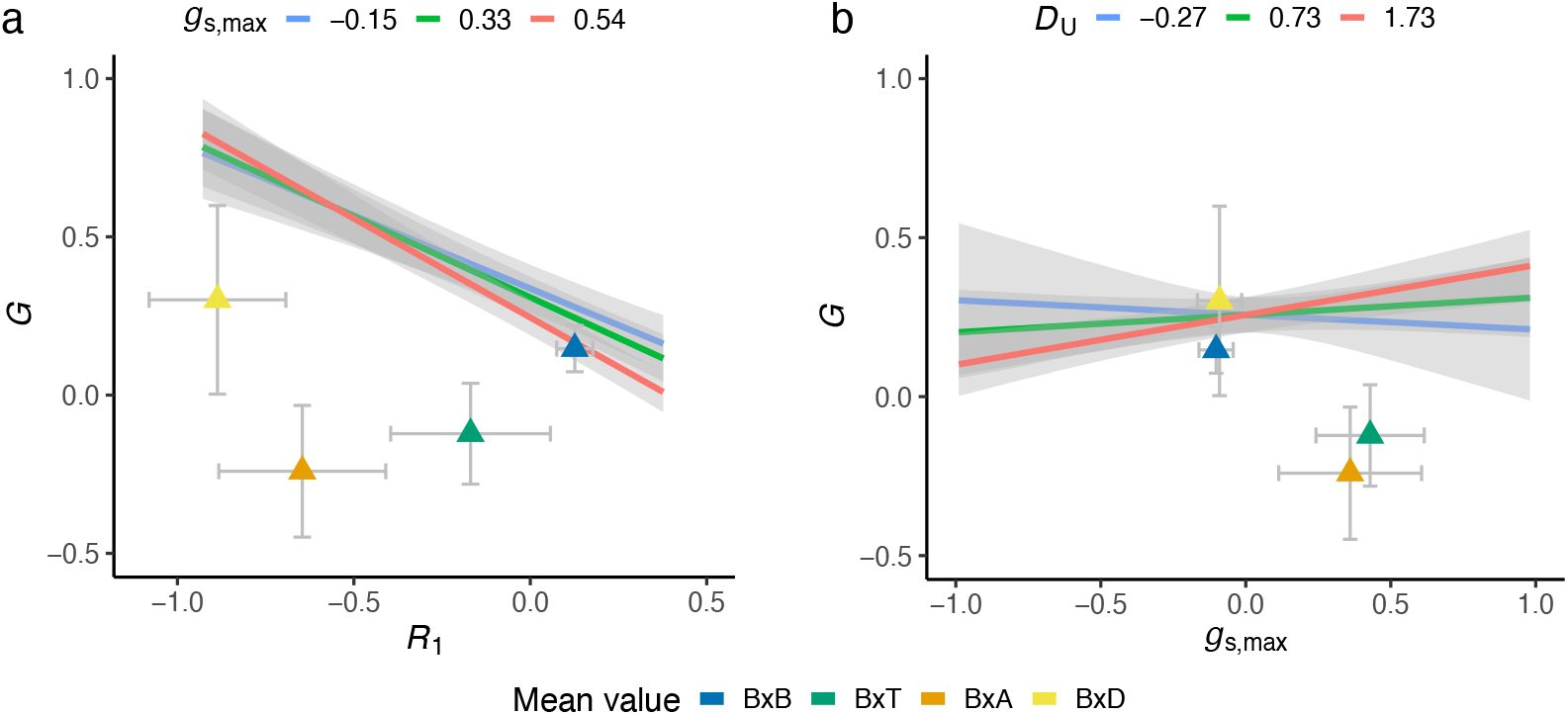
Contrasting growth-resistance and growth-gas exchange strategies are revealed between hybrid sets (model output from Eq. 12). The effects of resistance on growth are dependent on its interaction with *g*_s,max_ (a). Likewise, the effects of *g*_s,max_ on growth are dependent on its interaction with upper stomatal density (*D*_U_) (b). Mean BLUP value of hybrid sets and their 95% confidence intervals are plotted as triangles.

## IV. DISCUSSION

We conducted a quantitative genetic study in a set of *Populus* hybrids segregating for variation of disease resistance, stomatal patterning, and ecophysiological traits. Our motivations were to understand how hybridization changes trait distributions and heritable variation, and to identify correlations between stomatal and ecophysiology traits to disease resistance. We were particularly interested in identifying potential trade-offs and selection conflicts between pairs of traits that were masked in unadmixed populations but visible in hybrids. Our final motivation was to determine if we could identify competing growth strategies present in hybrid genotypes informing us about the evolution of growth strategies in natural hybrid zones, and potentially growth strategy evolution in each parental species. Our results clearly indicate hybridization, which we documented at three levels, has an important influence on trait values and the magnitude of their variances. Upper stomatal and ratio traits were correlated to variation of disease resistance, and we observed negative correlations between *D*_U_ and AR to resistance. After accounting for the effect of *D*_U_ variation (and all other predictors in the multi-response model), shifting stomatal area to the upper surface (i.e. increasing AR) or increasing the size of stomata on the upper surface decreased resistance. We observed a negative relationship between stomatal cover (*f*_S_) and resistance, and after accounting for *f*_S_, increasing *g*_s,max_ increases resistance. Trade-offs between *g*_s,max_ and *G* masked in the BxB set were made visible through hybridization in other sets. Likewise, the trade-off between *G* and R1 reversed sign in one hybrid set that was negative in the unadmixed *P. balsamifera* set. It is clear that hybridization is a useful tool for revealing trade-offs and studying the integration of sets of traits. We observed contrasting growth strategies along the growth-defense spectrum that were fined tuned by variation of *g*_s,max_ and *D*_U_. Indirectly, our results suggest selection is likely able to efficiently act on traits with high heritability and also correlated to disease resistance. Furthermore, the expression of trade-offs is dependent on the genetic background and environmental context. Finally, these results suggest the evolution of competing growth-defense strategies and their mis-alignment in hybrid zones may reinforce species boundaries, and that maladaptive genotypic variation observed in hybrids may have a phylogenetic context.

A question raised by these results is whether selection acting in these hybrid populations could sufficiently purge maladaptive genotypic variation linked to disease susceptibility? Our results suggest the answer is yes, resistance could be increased from selection for traits highly correlated to disease resistance, which, additionally, had large absolute value of scalar products to the proportion of *P. balsamifera* ancestry. Many of the traits with high absolute scalar product values also had high broad-sense heritabilities. For example, *D*_U_ and AR are two traits significantly correlated to disease resistance (regression coefficient = −9.67, −7.26, respectively) with large scalar products to *P. balsamifera* ancestry (PLS scalar product = −0.529, −0.524, respectively) and moderately high broad-sense heritability (*H*^2^ = 0.73, 0.4, respectively). If selection for increased disease resistance were to occur that targeted *D*_U_, AR, or traits with high co-variances to those two traits, populations similar to the ones we describe are likely to leave descendants with increased resistance. Similarly, any of the stomatal traits in the PLS blocks 1, 3, or 4 (Fig. 3) are likely good candidates to respond to selection for increased resistance.

The presence of amphistomy (having stomata on both leaf surfaces) as a result of hybridization raises interesting evolutionary consequences. Theoretical models of the benefits and costs of stomatal distributions indicate the increased efficiency of photosynthesis under amphistomy should lead to more species organizing their stomata on both surfaces (Muir, 2015). Yet, hypostomy (stomata only on the lower surface) predominates, particularly in trees and shrubs. Models which incorporate costs of amphistomy indicate a narrow range of optima, with few intermediate values of stomatal ratio, should predominate (Muir, 2015). The results from this study support theoretical conclusions that the costs of amphistomy are sufficiently high to constrain the available trait space in which plants evolve, particularly when pathogen pressure is high. While amphistomy is rare in *P. balsamifera*, it is common in *P. trichocarpa*, where northern populations have evolved increased stomatal ratio, possibly as a response to fine-tuning the growth-defense trade-off (McKown et al., 2014; McKown et al., 2019). We observed stomata on the upper leaf surfaces of *P. angustifolia* hybrids, possibly indicating selection for locally adapted stomatal phenotypes in the parental species’ populations. Further simulation work by Muir concludes that greater stomatal size or density increases the probability of pathogen colonization, and the effect is most pronounced when the fraction of leaf surface covered by stomata is low. Our results support Muir’s conclusions, as we demonstrate a pronounced decrease in resistance when stomatal densities are low and stomatal size is shifted to the upper leaf surface (Fig. 4b).

Evolution of growth strategies within species has been documented in other taxa and theory around growth-strategy evolution was important in early work that conceptually defined trade-offs (Schluter et al., 1991). Given sufficient selection from pathogens and heritable variation, plant species are likely to evolve a growth-defense optimum maximizing their fitness based on the likelihood of pathogen exposure, physiological severity of the disease, and the cost of mounting a defense (Obeso, 2002). The growth-defense optimum can be locally adapted, and even change between populations within a species (e.g. *P. trichocarpa,* McKown et al., 2014). When hybrids are formed, misaligned growth-defense strategies and the breakdown of phenotypic integration can negatively impact fitness through outbreeding depression (Goldberg et al., 2005), perhaps even in the presence of heterosis, as suggested by the decreased resistance in F1 BxD hybrids. These data may indirectly inform us about the evolution of growth strategies in hybrid zones or of the parental species themselves. Our data suggest *P. angustifolia* hybrids possess a fast-growing/low defense growth strategy, paired with higher *g*_s,max_ and *D*_U_ (Fig. 5), consistent with other reports from this species (e.g, Kaluthota et al., 2015). The observed growth rate of BxA hybrids is well below the predicted growth rate of the model, possibly as a result of the increased infection by *Melampsora* reducing growth. We may expect the observed growth rate of BxA hybirds to be closer to it’s prediction in an environment free of disease. *P. trichocarpa* hybrids are shifted along the spectrum, and have lower growth and higher resistance, paired with high *g*_s,max_ and higher *D*_U_, also consistent with reports from this species (e.g. McKown et al., 2014; McKown et al., 2019). *P. balsamfiera* genotypes appear to have evolved towards the lower growth/higher defense strategy and have lower *g*_s,max_ and almost a complete lack of upper stomata. Hybrids with P *deltoides* were all F_1_’s and our interpretation of these results is likely biased by heterosis, although they appear to have evolved towards the fast growth/low defense end of the spectrum and all BxD hybrids bear stomata on the upper leaf surface. Inferring growth strategies for parental species from hybrids is difficult. An experiment growing unadmixed genotypes of each species collected from their core ranges in a common environment would yield results free from the effects of admixture

Our results indicate an important role for resistance, *g*_s,max_ and *D*_U_ in fine tuning growth strategies. Our models predict that genotypes at the slow-growth/high-resistance end of the spectrum can increase their growth by decreasing *g*_s,max_, possibly suggesting that disease susceptibility is more costly to growth than a reduction in gas exchange (Fig. 5a). Growth can be finetuned by variation of *g*_s,max_ and *D*_U_, where optimal growth can be achieved with low *g*_s,max_ and *D*_U_ values, or, in contrast, high values of *g*_s,max_ and *D*_U_ (Fig. 5b). The interaction of *D*_U_ and *g*_s,max_ to increase growth at high values demonstrates a potential benefit to carrying stomata on the upper leaf surface, despite the higher risk of disease.

Hybridization appears to be an effective tool for disrupting phenotypic integration and introducing genotypic variance into a population. The conditional effects analyses on the centered and rescaled data demonstrated well how variance is introduced into a breeding population. In each of the three levels of hybridization we investigated, variance decreased towards the unadmixed population. It seems likely in populations similar to ours that selection would act at the larger scale of dozens of Mb to large portions of chromosomes, rather than soft selective sweeps acting on individual genes or causal variants. Although linkage decays rapidly in wind-pollinated, out-crossing species such as poplars (Tuskan et al., 2006), and adaptive introgression has been documented in these taxa before (e.g. Suarez-Gonzalez et al., 2018; Chhatre et al., 2018), the overwhelming fate of the non-*P. balsamifera* genetic material is mostly likely extirpation, even under mildly negative or purifying selection.

The magnitude of trade-offs and selection conflicts are not constant within our populations and are dependent on the genetic background, the environment, and the magnitude of genotype by environment (GxE). Trade-offs and selection conflicts may generally be subject to these interactions and visible in some circumstances, but not others. Although the data we present give us scant opportunity to determine the role of plasticity in revealing trade-offs and conflicts, the resistance data was measured in two years. We observed different estimates of *H*^2^ values and their variances between years, and the sizes of trade-offs were different within hybrid sets. A reversal of the growth-resistance trade-off was observed in the BxT hybrids. While the selection conflicts we observed were not significant, they reversed sign and magnitude between *R*_1_ and *R*_2_. These data indirectly support the idea that plasticity has an important evolutionary role for revealing or masking trade-offs and selection conflicts. Metaanalyses of plasticity have found adaptive plasticity to be less common than non-plastic modes of adaptation (Palacio-López et al., 2015), but the role of adaptive plasticity in maintaining fitness in a hybrid zones is less understood.

Collecting hybrids from crosses between multiple species within *Populus* allows us to indirectly infer the effect of phylogenetic distance of the parental species on trait variance, evolutionary, and ecological effects. We observed decreasing resistance with increasing phylogentic distance of the non-*balsamifera* parent after accounting for hybrid set (Fig. 2b). Resistance was restored with backcrossing into *P. balsamifera* advanced generation hybrids (Fig. S5). Increased disease in hybrid populations has been speculated to be an important ecological and evolutionary factor in maintaining species barriers (Bever et al., 2015). The increased disease we observed may be a common feature of hybrid zones, and indeed, hybrid zones may even provide refuge for pathogens and pests (Whitham, 1989). An example from a tri-hybrid zone in Alberta, Canada documents naturally formed F_1_ *P. balsamifera* x *angustifolia* hybrids transgressively segregating for the number of galls per tree and back-crosses into *P. balsamifera* were even more susceptible to gall forming pests and had higher resistance variance than F1 or unadmixed parental genotypes (Floate et al., 2016). These trends suggest an important role for pathogen associated selection in maintaining species barriers in *Populus*.

## V. CONCLUSIONS

We investigated the effects of hybridization on disease resistance and correlated stomatal and ecophysiological traits. We have demonstrated the effects of hybrdization on trait variance at multiple scales, and shown how hybridization can reveal trade-offs and potential selection conflicts when integrated modules of traits, likely adapted in either parental species, are combined in admixed populations. Misalignment of growth-defense strategies results in decreased disease resistance and maladapive phenotypic distributions. We are able to better understand how pathogen-associated selection can constrain stomatal trait distributions in admixed populations. These results demonstrate the important evolutionary and ecological effects of hybridization in plant-pathogen interactions. Future research in this system will focus on using admixture mapping to identify genomic regions which underlie the disease resistance observed in *P. balsamifera* and *P. trichocarpa*. Understanding the core growth-defense strategies that have evolved in each species will allow ecologists to place their findings of disease ecology in hybrid zones into a wider context, and should be undertaken by future researchers.

## VI. DATA ACCESSIBILITY

Cuticle micrographs are deposited on Dryad (doi:10.5061/dryad.kh2gv5f). Raw sequence reads are available on NCBI SRA (accession # SRP070954). NewHybrids filial call probabilities are provided in Supplementary Information S4a-c. Data used for this study are provided in the appendix 2.

## X. SUPPORTING INFORMATION

**Table S3:**
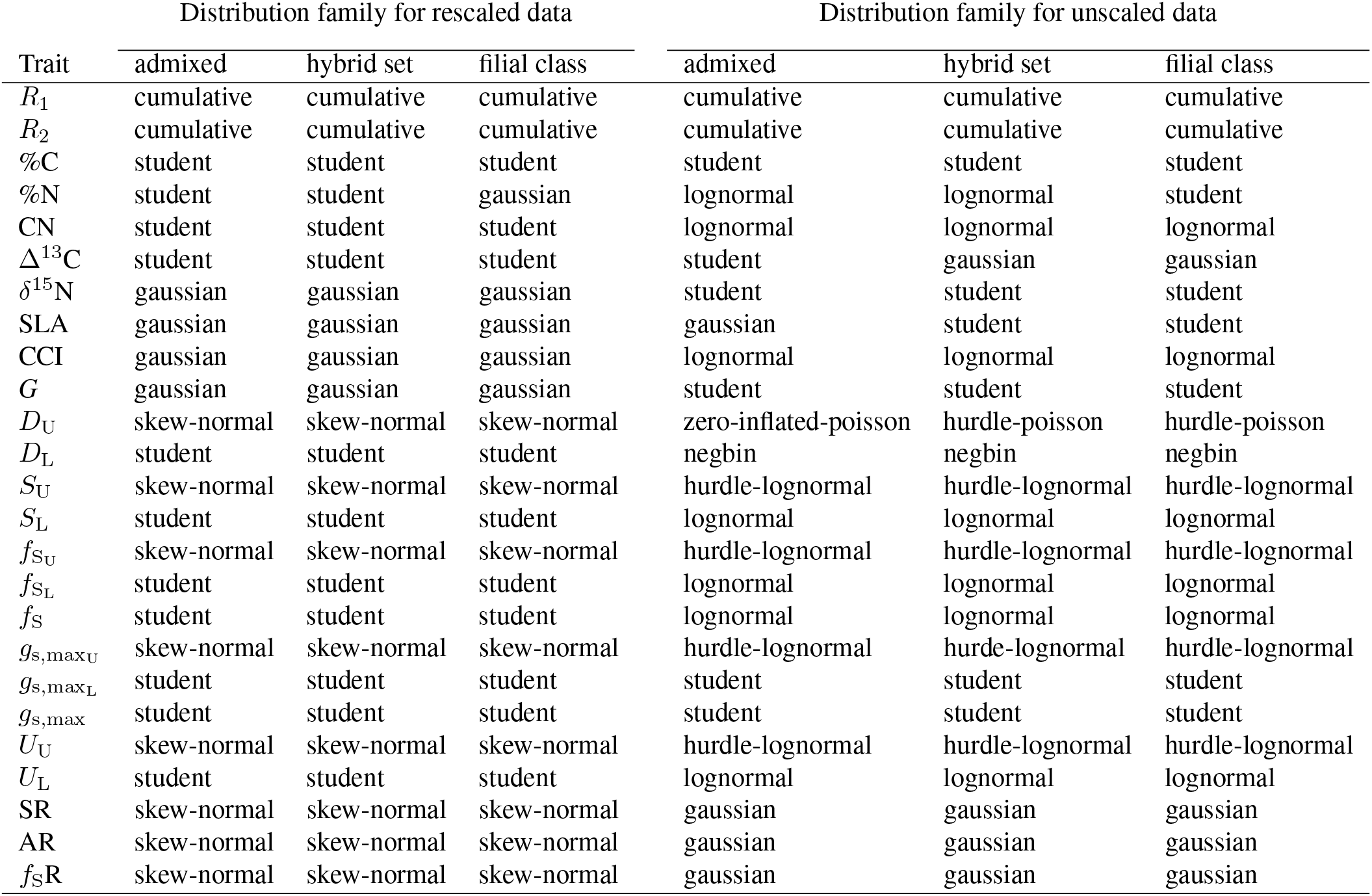
Distribution families used in Bayesian multi-level models for rescaled and unsealed data. Models were fit with brms (Bürkner, 2017).

**Table S4:** Multi-response models tested with LOO cross validation. The responses in each model were *R*_1_ and *R*_2_. Models were fit and evaluated with brms (Bürkner, 2017). ΔELPD is the expected log point wise predictive density for a new dataset, a measure of difference of the fit of the best model to itself. *A, Z,* and *I* are the matrix of garden xy coordinates, expected ancestry proportion, and the random effects of individual, respectively.

| Model ID | Model | $\Delta\text{ELPD}$ |
| --- | --- | --- |
| 1 | $D_U + D_L + S_U + S_L + fS + g_{s,\max} + SR + AR + G + \Delta^{13}C + \delta^{15}N + CN + SLA + CCI + A + Z + I$ | 0.0 |
| 2 | $D_U + D_L + S_U + S_L + fS + g_{s,\max} + SR + AR + fsR + G + \Delta^{13}C + \delta^{15}N + CN + SLA + CCI + A + Z + I$ | -4.3 |
| 3 | $fS + g_{s,\max} + SR + AR + fsR + G + \Delta^{13}C + \delta^{15}N + CN + SLA + CCI + A + Z + I$ | -21.8 |
| 4 | $fS + g_{s,\max} + SR + AR + G + \Delta^{13}C + \delta^{15}N + CN + SLA + CCI + A + Z + I$ | -23.0 |

**Figure S1:**
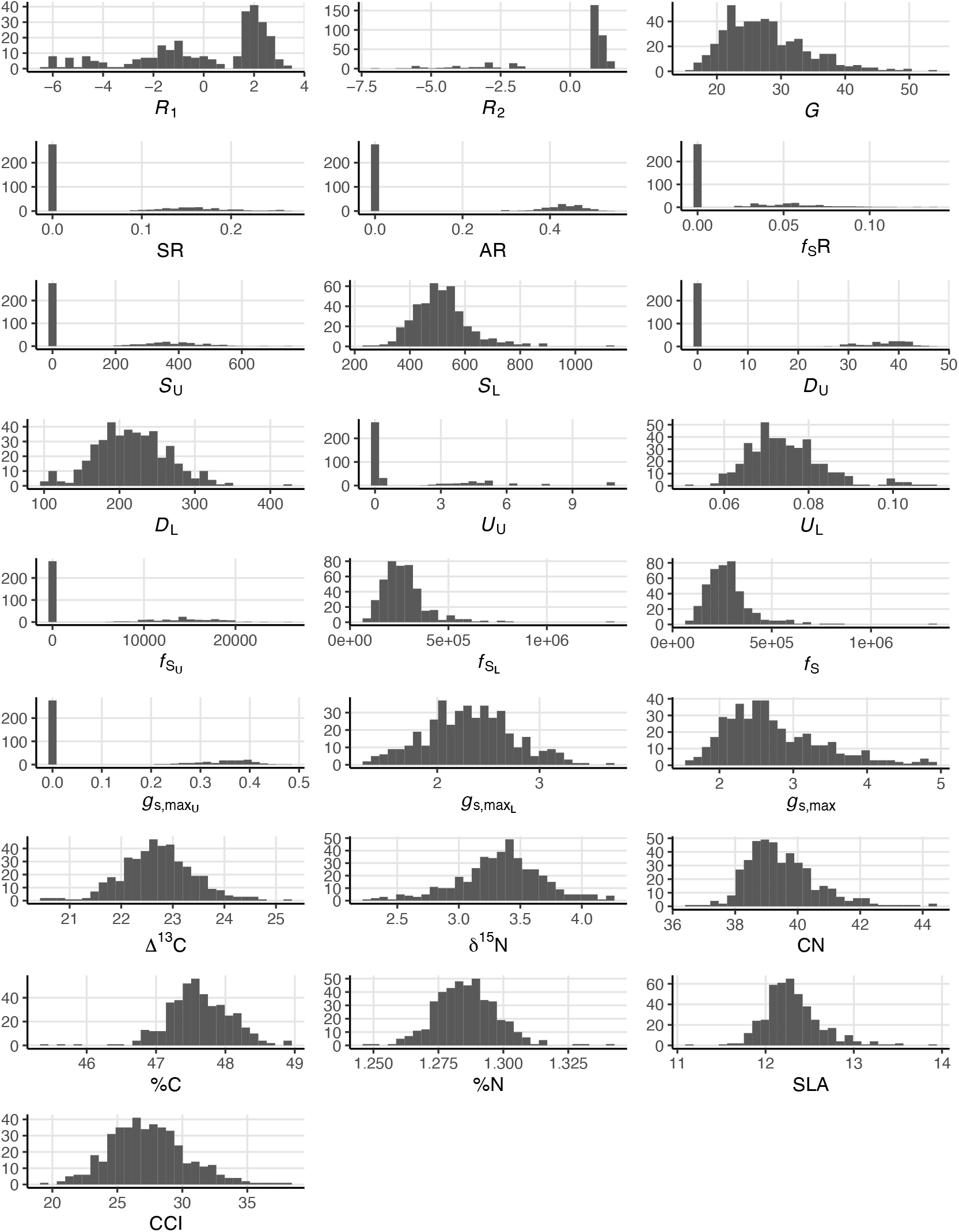
Distribution of traits modelled as BLUPs (*R*_1_ and *R*_2_) and mBLUPs (remaining traits). See Table 1 for trait abbreviations and units.

**Figure S2:**
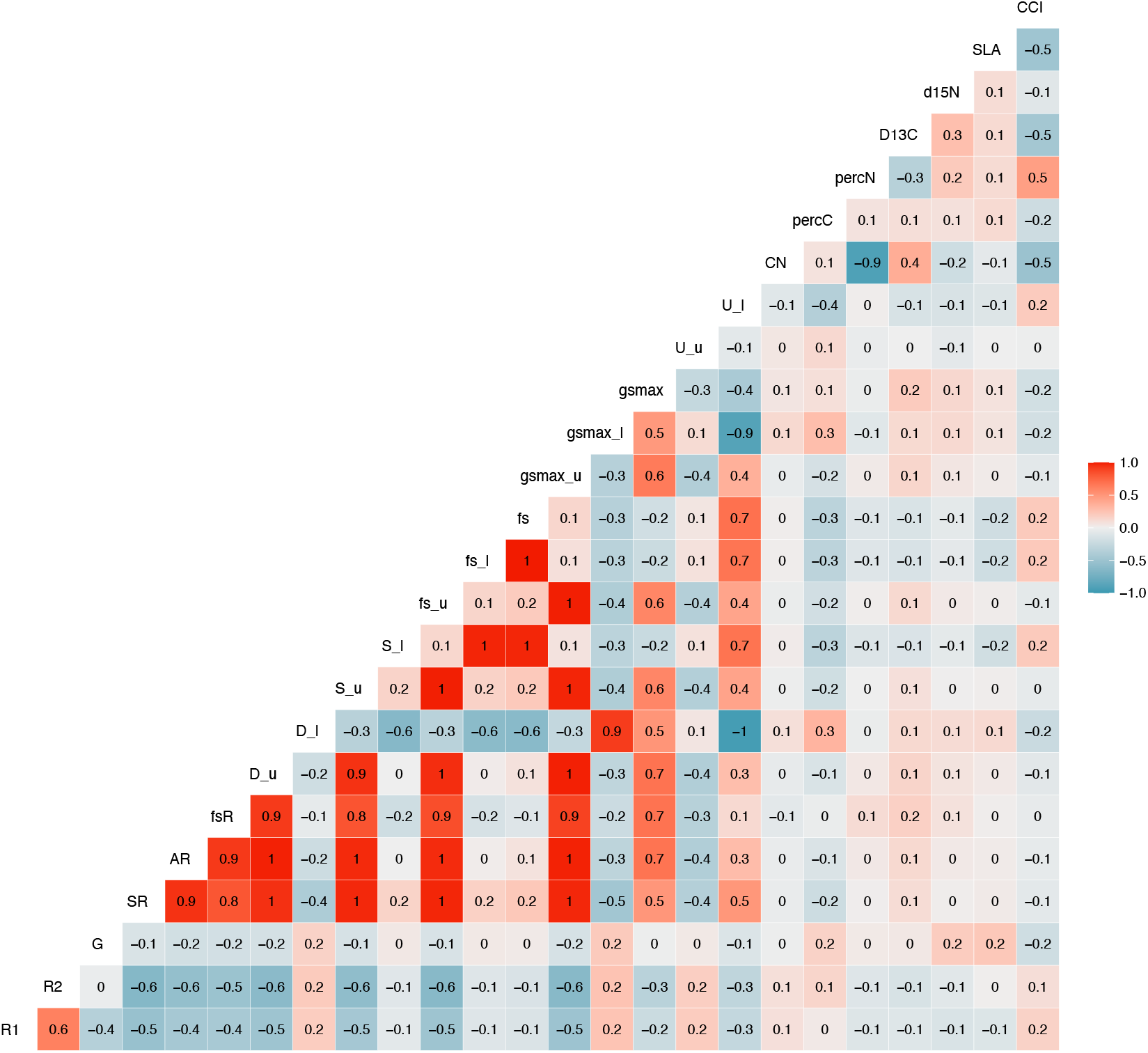
Pairwise Pearson’s correlation coefficients. See Table 1 for trait abbreviations and units.

**Figure S3:**
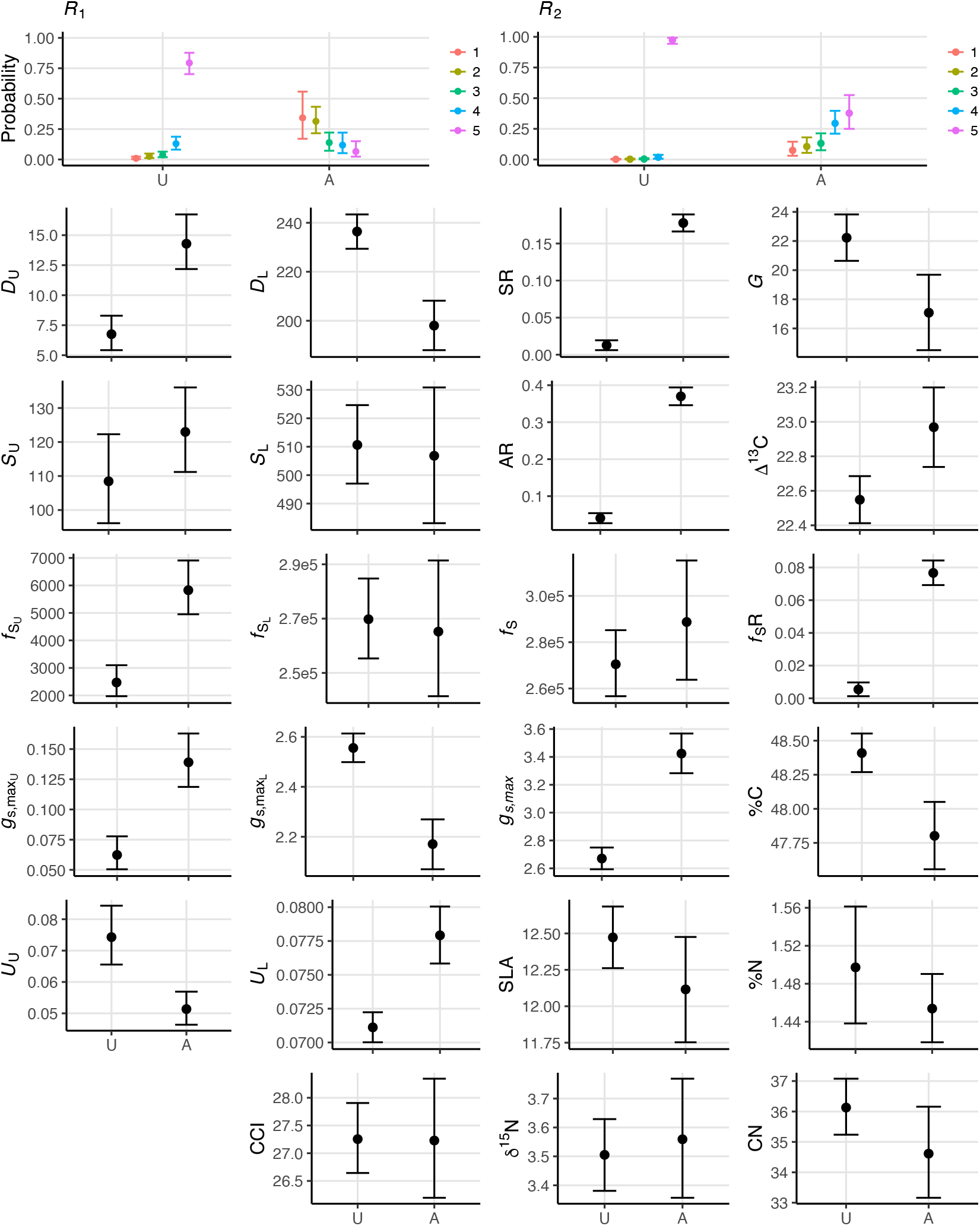
Conditional effects of admixture status on trait variation. See Table 1 for trait definitions. Abbreviations: U, unadmixed; A, admixed.

**Figure S4:**
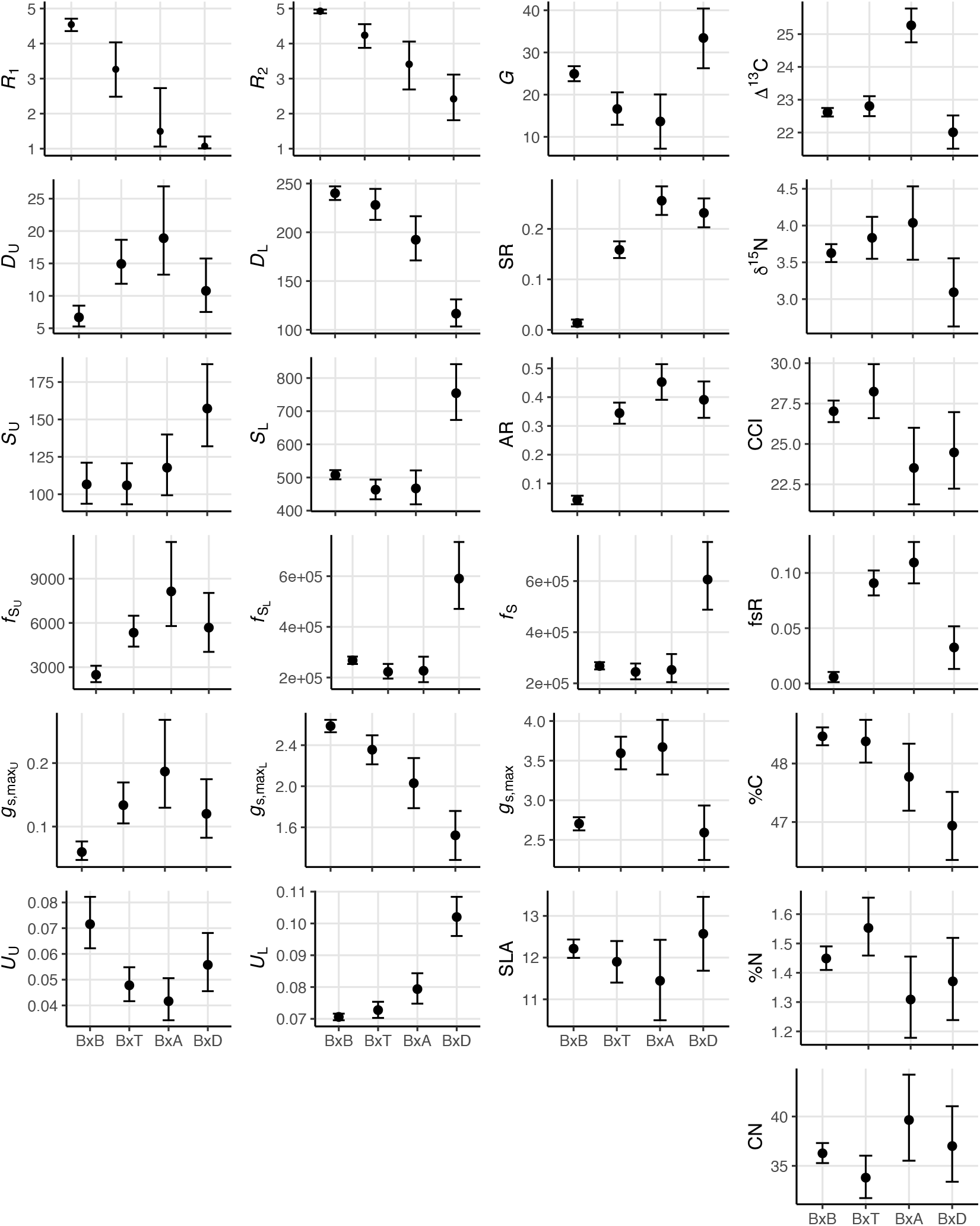
Conditional effects of hybrid set on trait variation. See Table 1 for trait definitions.

**Figure S5:**
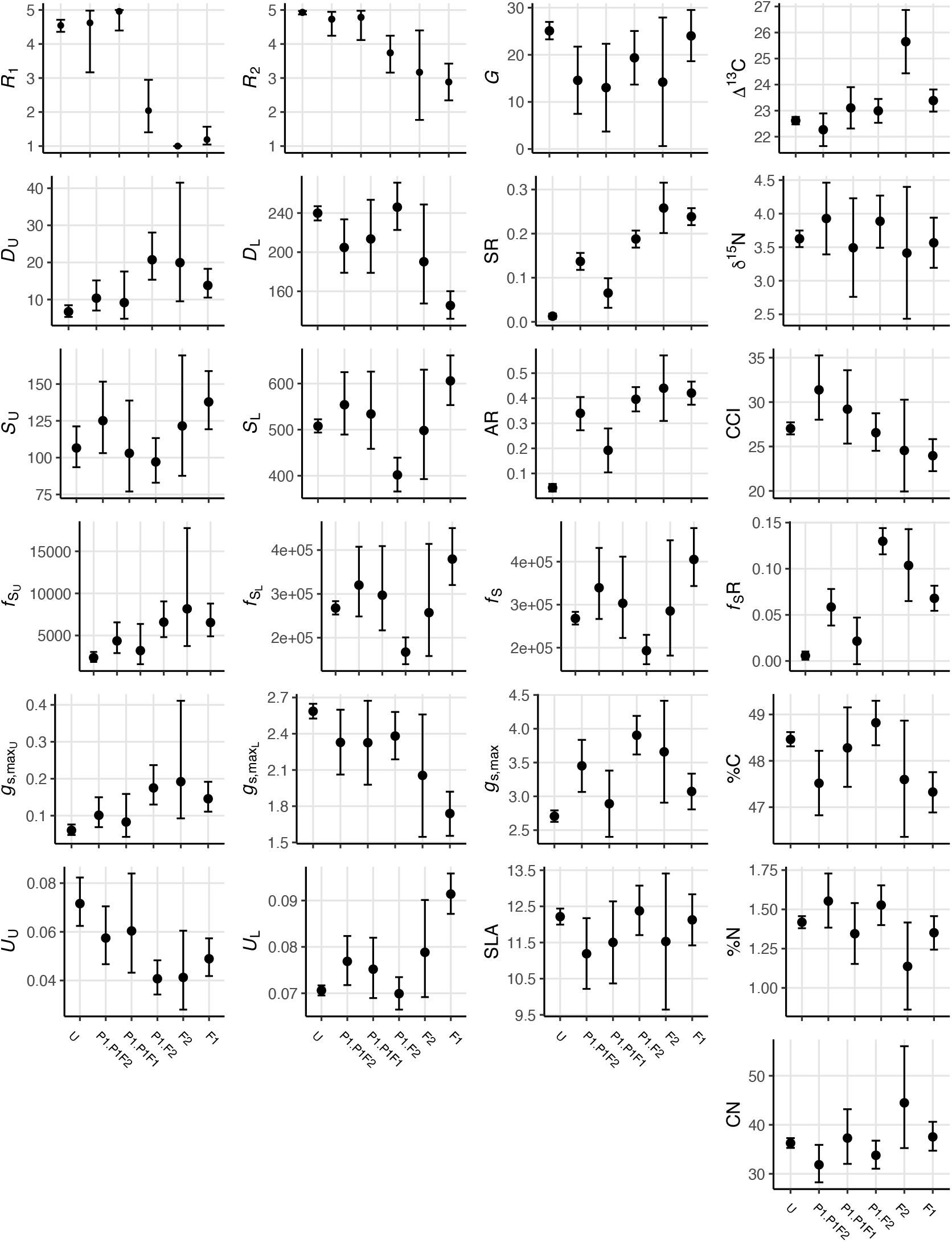
Variation of unscaled traits by filial generation. See Table 1 for trait defintions; abbreviations; U, unadmixed.

**Figure S6:**
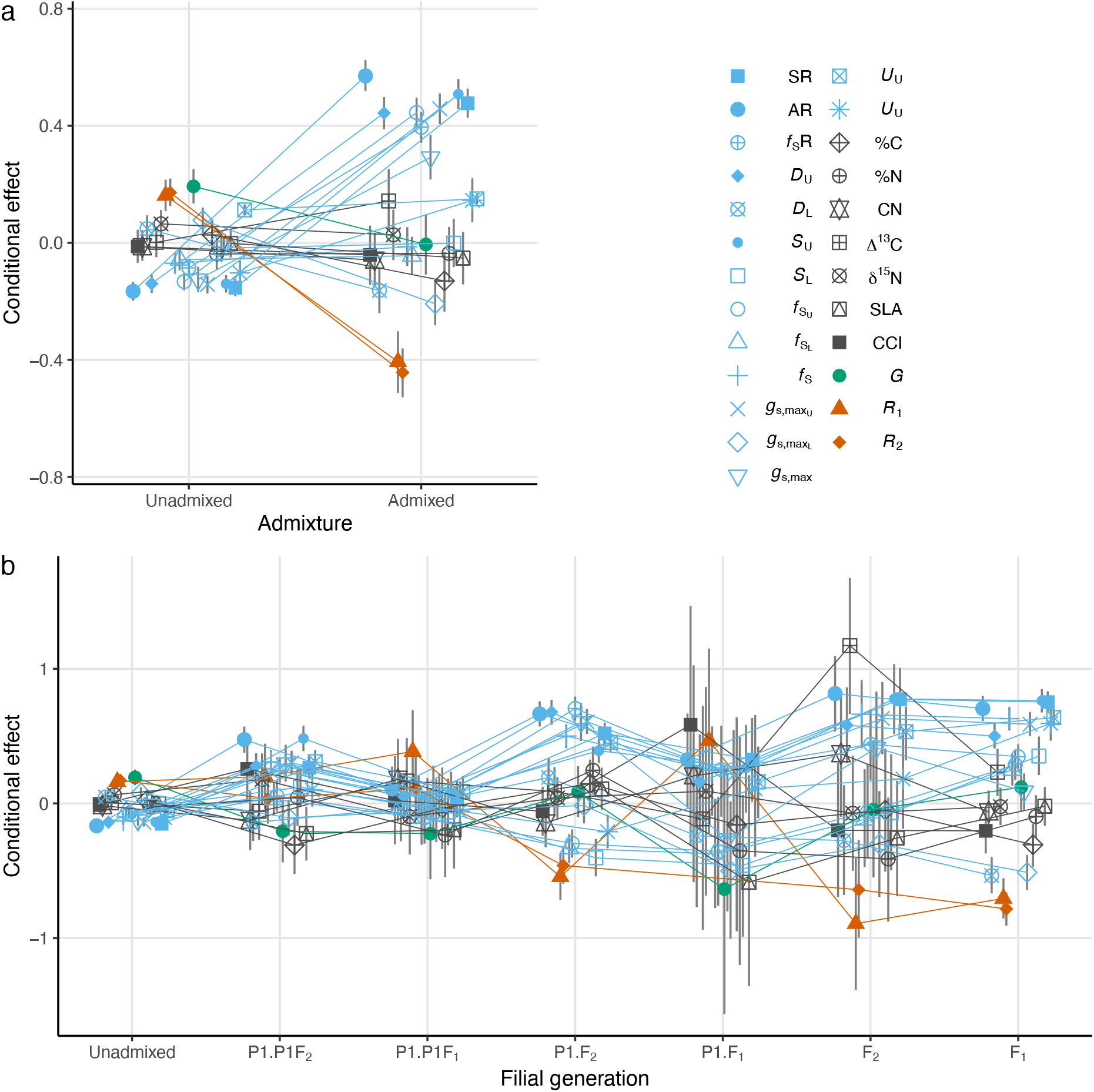
Variation of rescaled traits by admixture status and filial generation (Eq. 8). See Table 1 for trait abbreviations.

## REFERENCES

Allen, E, B Hazen, H Hoch, Y Kwon, G Leinhos, R Staples, M Stumpf, B Terhune, et al. (1991). “Appressorium formation in response to topographical signals by 27 rust species.” In: Phytopathology 81.3, pp. 323–331.

Anderson, E and E Thompson (2002). “A model-based method for identifying species hybrids using multilocus genetic data”. In: Genetics 160.3, pp. 1217–1229.

Bever, JD, SA Mangan, and HM Alexander (2015). “Maintenance of plant species diversity by pathogens”. In: Annual review of ecology, evolution, and systematics 46, pp. 305–325.

Browning, BL, Y Zhou, and SR Browning (2018). “A one-penny imputed genome from next-generation reference panels”. In: The American Journal of Human Genetics 103.3, pp. 338–348.

Bruns, EL, J Antonovics, and M Hood (2018). “Is there a disease-free halo at species range limits? The co-distribution of anther-smut disease and its host species”. In: Journal of Ecology.

Bürkner, PC (2017). “brms: An R Package for Bayesian Multilevel Models Using Stan”. In: Journal of Statistical Software 80.1, pp. 1–28.

Chandran, D, J Rickert, Y Huang, MA Steinwand, SK Marr, and MC Wildermuth (2014). “Atypical E2F transcriptional repressor DEL1 acts at the intersection of plant growth and immunity by controlling the hormone salicylic acid”. In: Cell host & microbe 15.4, pp. 506–513.

Chhatre, VE, LM Evans, SP DiFazio, and SR Keller (2018). “Adaptive introgression and maintenance of a trispecies hybrid complex in range-edge populations of *Populus*”. In: Molecular ecology.

Chhatre, VE, KC Fetter, AV Gougherty, MC Fitzpatrick, RY Soolanayakanahally, RS Zalensy, and SR Keller (2019). “Climatic niche predicts the landscape structure of locally adaptive standing genetic variation”. In: BioRxiv, p. 817411.

Christe, C, KN Stölting, L Bresadola, B Fussi, B Heinze, D Wegmann, and C Lexer (2016). “Selection against recombinant hybrids maintains reproductive isolation in hybridizing *Populus* species despite F1 fertility and recurrent gene flow”. In: Molecular Ecology 25.11, pp. 2482–2498.

Danecek, P, A Auton, G Abecasis, CA Albers, E Banks, MA DePristo, RE Handsaker, G Lunter, GT Marth, ST Sherry, et al. (2011). “The variant call format and VCFtools”. In: Bioinformatics 27.15, pp. 2156–2158.

Dangl, JL and JD Jones (2001). “Plant pathogens and integrated defence responses to infection”. In: nature 411.6839, p. 826.

DiFazio, SP, GT Slavov, and CP Joshi (2011). “Populus: a premier pioneer system for plant genomics”. In: Enfield, NH: Science Publishers, pp. 1–28.

Eckenwalder, JE (1996). “Systematics and evolution of Populus”. In: Biology of Populus and its implications for management and conservation. Ed. by R Stettler, T Bradshaw, P Heilman, and T Hinckley. Ottawa: NRC Research Press. Chap. 1, pp. 7–32.

Elshire, RJ, JC Glaubitz, Q Sun, JA Poland, K Kawamoto, ES Buckler, and SE Mitchell (2011). “A robust, simple genotyping-by-sequencing (GBS) approach for high diversity species”. In: PloS one 6.5, e19379.

Evans, LM, GT Slavov, E Rodgers-Melnick, J Martin, P Ranjan, W Muchero, AM Brunner, W Schackwitz, L Gunter, JG Chen, et al. (2014). “Population genomics of *Populus trichocarpa* identifies signatures of selection and adaptive trait associations”. In: Nature genetics 46.10, p. 1089.

Farquhar, GD, MH O’Leary, and JA Berry (1982). “On the relationship between carbon isotope discrimination and the intercellular carbon dioxide concentration in leaves”. In: Functional Plant Biology 9.2, pp. 121–137.

Feau, N, DL Joly, and RC Hamelin (2007). “Poplar leaf rusts: model pathogens for a model tree”. In: Botany 85.12, pp. 1127–1135.

Fetter, KC, S Eberhardt, RS Barclay, S Wing, and SR Keller (2019). “StomataCounter: a neural network for automatic stomata identification and counting”. In: New Phytologist 223.3, pp. 1671–1681.

Floate, KD, J Godbout, MK Lau, N Isabel, and TG Whitham (2016). “Plant-herbivore interactions in a trispecific hybrid swarm of *Populus*: assessing support for hypotheses of hybrid bridges, evolutionary novelty and genetic similarity”. In: New Phytologist 209.2, pp. 832–844.

Gelman, A (2008). “Scaling regression inputs by dividing by two standard deviations”. In: Statistics in medicine 27.15, pp. 2865–2873.

Glaubitz, JC, TM Casstevens, F Lu, J Harriman, RJ Elshire, Q Sun, and ES Buckler (2014). “TASSEL-GBS: a high capacity genotyping by sequencing analysis pipeline”. In: PloS one 9.2, e90346.

Goldberg, TL, EC Grant, KR Inendino, TW Kassler, JE Claussen, and DP Philipp (2005). “Increased infectious disease susceptibility resulting from outbreeding depression”. In: Conservation Biology 19.2, pp. 455–462.

Gonzales-Vigil, E, CA Hefer, ME von Loessl, J La Mantia, and SD Mansfield (2017). “Exploiting natural variation to uncover an alkene biosynthetic enzyme in poplar”. In: The Plant Cell, tpc–00338.

Goulet, BE, F Roda, and R Hopkins (2017). “Hybridization in plants: old ideas, new techniques”. In: Plant physiology 173.1, pp. 65–78.

Kaluthota, S, DW Pearce, LM Evans, MG Letts, TG Whitham, and SB Rood (2015). “Higher photosynthetic capacity from higher latitude: foliar characteristics and gas exchange of southern, central and northern populations of Populus angustifolia”. In: Tree physiology 35.9, pp. 936–948.

Kliebenstein, DJ (2016). “False idolatry of the mythical growth versus immunity tradeoff in molecular systems plant pathology”. In: Physiological and Molecular Plant Pathology 95, pp. 55–59.

La Mantia, J, J Klápšte, YA El-Kassaby, S Azam, RD Guy, CJ Douglas, SD Mansfield, and R Hamelin (2013). “Association analysis identifies Melampsora× columbiana poplar leaf rust resistance SNPs”. In: PLoS One 8.11, e78423.

Lê, S, J Josse, F Husson, et al. (2008). “FactoMineR: an R package for multivariate analysis”. In: Journal of statistical software 25.1, pp. 1–18.

Li, H and R Durbin (2009). “Fast and accurate short read alignment with Burrows-Wheeler transform”. In: bioinformatics 25.14, pp. 1754–1760.

McKown, AD, RD Guy, L Quamme, J Klápšte, J La Mantia, C Constabel, YA El-Kassaby, RC Hamelin, M Zifkin, and M Azam (2014). “Association genetics, geography and ecophysiology link stomatal patterning in *Populus trichocarpa* with carbon gain and disease resistance trade-offs”. In: Molecular ecology 23.23, pp. 5771–5790.

McKown, AD, J Klápště, RD Guy, OR Corea, S Fritsche, J Ehlting, YA El-Kassaby, and SD Mansfield (2019). “A role for SPEECHLESS in the integration of leaf stomatal patterning with the growth vs disease trade-off in poplar.” In: The New phytologist.

Melotto, M, W Underwood, J Koczan, K Nomura, and SY He (2006). “Plant stomata function in innate immunity against bacterial invasion”. In: Cell 126.5, pp. 969–980.

Messina, FJ, SL Durham, JH Richards, and DE McArthur (2002). “Trade-off between plant growth and defense? A comparison of sagebrush populations”. In: Oecologia 131.1, pp. 43–51.

Muir, CD (2020). “A stomatal model of anatomical tradeoffs between gas exchange and pathogen colonization”. In: bioRxiv, p. 871228.

Muir, CD (2015). “Making pore choices: repeated regime shifts in stomatal ratio”. In: Proceedings of the Royal Society B: Biological Sciences 282.1813, p. 20151498.

Obeso, JR (2002). “The costs of reproduction in plants”. In: New Phytologist 155.3, pp. 321–348.

Palacio-López, K, B Beckage, S Scheiner, and J Molofsky (2015). “The ubiquity of phenotypic plasticity in plants: a synthesis”. In: Ecology and evolution 5.16, pp. 3389–3400.

Roche, B and R Fritz (1998). “Effects of host plant hybridization on resistance to willow leaf rust caused by *Melampsora* sp.” In: European Journal of Forest Pathology 28.4, pp. 259–270.

Rohart, F, B Gautier, A Singh, and KA Lê Cao (2017). “mixOmics: An R package for ‘omics feature selection and multiple data integration”. In: PLoS computational biology 13.11, e1005752.

Sack, Land TN Buckley (2016). “The developmental basis of stomatal density and flux”. In: Plant Physiology 171.4, pp. 23582363.

Schluter, D, TD Price, and L Rowe (1991). “Conflicting selection pressures and life history trade-offs”. In: Proc. R. Soc. Lond. B 246.1315, pp. 11–17.

Schneider, CA, WS Rasband, and KW Eliceiri (2012). “NIH Image to ImageJ: 25 years of image analysis”. In: Nature methods 9.7, p. 671.

Suarez-Gonzalez, A, C Lexer, and QC Cronk (2018). “Adaptive introgression: a plant perspective”. In: Biology letters 14.3, p. 20170688.

Tian, D, M Traw, J Chen, M Kreitman, and J Bergelson (2003). “Fitness costs of R-gene-mediated resistance in *Arabidopsis thaliana*”. In: Nature 423.6935, p. 74.

Tuskan, GA et al. (2006). “The genome of black cottonwood, *Populus trichocarpa* (Torr. & Gray)”. In: science 313.5793, pp. 1596–1604.

Ullah, C, CJ Tsai, SB Unsicker, L Xue, M Reichelt, J Gershenzon, and A Hammerbacher (2018). “Salicylic acid activates poplar defense against the biotrophic rust fungus *Melampsora larici-populina* via increased biosynthesis of catechin and proanthocyanidins”. In: New Phytologist.

Van Kraayenoord, C, G Laundon, and A Spiers (1974). “Poplar rusts invade New Zealand.” In: Plant disease reporter 58.5, pp. 423–427.

White, J, B Vaughn, and S Michel (2011). “Stable Isotopic Composition of Atmospheric Carbon Dioxide (13C and 18O) from the NOAA ESRL Carbon Cycle Cooperative Global Air Sampling Network, 1990-2014, Version: 2015-10-26.” In: University of Colorado, Institute of Arctic and Alpine Research (INSTAAR).

Whitham, TG (1989). “Plant hybrid zones as sinks for pests”. In: Science, pp. 1490–1493.

